# Quantitative synapse analysis for cell-type specific connectomics

**DOI:** 10.1101/386912

**Authors:** Dika A. Kuljis, Khaled Zemoura, Cheryl A. Telmer, Jiseok Lee, Eunsol Park, Daniel S. Ackerman, Weifeng Xu, Simon C. Watkins, Don B. Arnold, Marcel P. Bruchez, Alison L. Barth

## Abstract

Anatomical methods for determining cell-type specific connectivity are essential to inspire and constrain our understanding of neural circuit function. We developed new genetically-encoded reagents for fluorescence-synapse labeling and connectivity analysis in brain tissue, using a fluorogen-activating protein (FAP)-or YFP-coupled, postsynaptically-localized neuroligin-1 targeting sequence (FAP/YFPpost). Sparse viral expression of FAP/YFPpost with the cell-filling, red fluorophore dTomato (dTom) enabled high-throughput, compartment-specific localization of synapses across diverse neuron types in mouse somatosensory cortex. High-resolution confocal image stacks of virally-transduced neurons were used for 3D reconstructions of postsynaptic cells and automated detection of synaptic puncta. We took advantage of the bright, far-red emission of FAPpost puncta for multichannel fluorescence alignment of dendrites, synapses, and presynaptic neurites to assess subtype-specific inhibitory connectivity onto L2 neocortical pyramidal (Pyr) neurons. Quantitative and compartment-specific comparisons show that PV inputs are the dominant source of inhibition at both the soma and across all dendritic branches examined and were particularly concentrated at the primary apical dendrite, a previously unrecognized compartment of L2 Pyr neurons. Our fluorescence-based synapse labeling reagents will facilitate large-scale and cell-type specific quantitation of changes in synaptic connectivity across development, learning, and disease states.

## 1 Introduction

The organization, number, and input identity of synapses onto a cell are critical determinants of neuronal activity. Although electrophysiological analyses of synaptic properties have provided a rich framework to build and test hypotheses about neural computations during sensation and behavior, these analyses cannot reveal broader principles of synaptic distribution across the neuron. Since alterations to synaptic function in select circuits and cell types are associated with autism, intellectual disability, psychiatric, and neurologic disease (Bayes et al., 2011; Sudhof, 2017), quantitative metrics about synaptic location, size, and input specificity are likely to provide key insights into how specific diseases alter neural circuits.

Electron-microscopy (EM) provides nanometer resolution for ultrastructural identification of synaptic contacts and has been employed for brain-area and cell-type quantitative analysis (Bock et al., 2011; Briggman et al., 2011; Chandrasekaran et al., 2015; Kim et al., 2014); however, EM is hampered by technical demands of sample preparation, imaging time, data storage, and labor-intensive analysis that make comparison across multiple individuals or conditions difficult. Recent studies have attempted to use EM for quantitative analysis of synapse organization between defined pre- and postsynaptic partners, but these computationally-intensive approaches are difficult to adopt and scale for broad use (Glausier et al., 2017; Kornfeld et al., 2017; Kubota et al., 2015; Vishwanathan et al., 2017). Fluorescence-based microscopy methods are an attractive alternative to EM, because light-microscopy facilitates faster acquisition of larger tissue volumes and enables use of spectrally distinct, genetically encoded fluorophores for discrimination of molecularly diverse cells and synapse types.

There has been great interest in developing tools and methodologies for synapse labeling using molecular, genetic, or histochemical techniques for light microscopy, including GFP-tagging synaptic molecules, GFP reconstitution across synaptic partners (GRASP) and array tomography (Chen et al., 2012; Fortin et al., 2014; Gross et al., 2013; Kim et al., 2011; Martell et al., 2016; Micheva and Smith, 2007; Villa et al., 2016a). Fluorescence-based, sparse labeling of post-synaptic neurons in intact brain tissue has been especially helpful in this regard, as it reduces the analysis bottleneck that arises from non-specific immunohistochemical labeling of synapses. However, high-throughput/volumetric synaptic analysis for individual neurons has not yet become routine, due to low signal-to-noise and synaptogenesis associated with overexpression of synaptic tags (El-Husseini et al., 2000; Kim et al., 2011; Martell et al., 2016).

We sought to develop molecular genetic tools for comprehensive fluorescence labeling of postsynaptic sites across an individual neuron, in a complex tissue environment. We employed two tags for synaptic labeling using either YFP or the fluorogen-activating protein (FAP), a modified antibody fragment that emits in the far red upon binding of a small molecule ligand, a derivative of malachite green (Szent-Gyorgyi et al., 2013). These two extracellular tags were targeted to postsynaptic sites using the transmembrane and cytoplasmic region of mouse neuroligin-1 (NL-1) (Kim et al., 2011). Sparse, virus-mediated coexpression of FAP/YFPpost with the cell-filling fluorophore dTomato (dTom) showed broad, punctate labeling across distinct cell compartments and different cell types. Aided by automated image analysis, we quantitatively evaluated the distribution of more than 140,000 synaptic puncta across four molecularly distinct neocortical neuron subtypes. Multichannel fluorescence imaging in tissue from parvalbumin (PV), somatostatin (SST), and vasoactive intestinal peptide (VIP) Cre-driver transgenic mice allowed us to quantify cell-type specific inputs to FAPpost puncta in L2 Pyr neurons. This quantitative analysis revealed that PV inputs dominated the soma and the synapse-dense 1° apical dendrite, and that PV inputs had a higher density than SST inputs across L2 dendrites. These studies initiate a new paradigm for high-throughput analysis of synapse organization in brain tissue during health and disease.

## 2 Materials and Methods

All experimental procedures were conducted in accordance with the NIH guidelines and were approved by the Institutional Animal Care and Use Committee at Carnegie Mellon University.

### 2.1 Construct Design

#### 2.1.1 FAPpost Cloning

To make the plasmid for packaging into AAV, post-mGRASP from Addgene (#34912 - paavCAG-post-mGRASP-2A-dTom) was modified by annealing oligos and inserting into BamHI and XhoI digested backbone to introduce an AgeI site (PostBamXhoF 5’GATCC CTT ACCGGT ATC TTA C and PostBamXhoR 5’ TCGAG TAA GAT ACCGGT AAG G). PCR was used to produce the Igkappa leader sequence, cmyc epitope and dL5** FAP (Szent-Gyorgyi et al., 2008; Szent-Gyorgyi et al., 2013; Telmer et al., 2015) for introduction into the BamHI and AgeI of the modified backbone (BamKappaF 5’ TATATA GGATCC ggcttggggatatccaccatgg and dL5AgeSfiR 5’ TATATA ACCGGT ACCTCC ggccagaccggccgc GGAGAG). The BamHI/HindIII fragment was moved to create pENN.AAV.hSyn.kappa.myc.dL5.POSTsyn.T2A.dTom.WPRE.BGH (Addgene FAPpost plasmid ID 105981). AAV1 serotype was produced by Penn Vector Core.

#### 2.1.2 Fl-YFPpost and fl-FAPpost Cloning

For Cre-inducible expression, the kappa.myc.dL5.POSTsyn.T2A.dTom region was PCR amplified with primers containing BsrG1 and KpnI restriction sites (partial KpnI digestion was required) and ligated into digested pAAV-FLEX (fl; generous gift from Oliver Schluter) to produce pAAV-FLEX-hSyn-kappa-myc-dL5-POSTsyn-T2A.dTom-WPRESV40. PCR amplification was used to generate the SYFP2 (YFP) (Kremers et al., 2006) coding fragment (iGEM BBa_K864100) that was then SfiI digested to replace the FAP in the pAAV-FLEX resulting in pAAV-FLEX-hSyn-kappa-myc-dL5-POSTsyn-T2A-dTom-WPRE-SV40 (Addgene fl-FAPpost plasmid ID 105982; Addgene fl-YFPpost plasmid ID 105983). Constructs were packaged into AAV1 and produced by Penn Vector Core.

#### 2.1.3 PSD-95 and Gephyrin FingRs

The open reading frame encoding Gephyrin or PSD-95 FingRs was fused to GFP and the CCR5 Zinc FingR DNA binding domain (Gross et al., 2013) and double floxed and inverted for Cre-dependent expression. Expression was driven by the CAG (beta-actin) promoter. AAV serotype 9 was generated at UPenn Vector Core (AAV9.CAG.DIO.GPHN.Fn-eGFP-ZF.hGH (4.98kb) and AAV9.CAG.DIO.PSD95.Fn-eGFP-ZF.hGH (4.98kb)).

### 2.2 Neuronal cultures and immunostaining

Primary neuronal cultures of cerebral cortex were prepared from C57BL6 newborn mice, aged P0/day of birth. Neurons were plated to a density of about 250,000 cells onto poly-L-lysine-coated coverslips. Neurons were kept in culture at 37 °C and 5% CO_2_ for 11–21 days. At day 3 neurons were infected with AAV FAPpost (Shi et al., 2017). Twelve days after infection, the neurons were rinsed with PBS and fixed for 10 minutes using 4% PFA/PBS at room temperature. After fixation, cells were again rinsed in PBS and permeabilized for 7 minutes using 0.2% Triton-X/PBS. The neurons were then rinsed in PBS and incubated overnight in a hydrated chamber at 4°C with an anti-PSD-95 (1:250; Synaptic Systems 124 011) or anti-gephyrin antibody (1:200, Synaptic Systems 147 011C3) diluted in PBS/10% NDS. The coverslips were then washed 3 times for 5 minutes with PBS followed by incubation in the dark for 45 minutes in a hydrated chamber with a secondary antibody (Alexa-Fluor 488-conjugated donkey anti-mouse; Jackson Immunoresearch 715-545-150) diluted1:800 in PBS/10% NDS and the fluorogenic malachite green dye derivative MG-tcarb (MG; 300nM in PBS; (Pratt et al., 2017)). Finally, the cultures were washed with PBS, dried, and mounted onto glass slides with Dako fluorescent mounting medium (Zemoura et al., 2016). Images were collected by laser Airyscan confocal microscopy (ZEISS Airyscan LSM 880). A z-stack of five to eight optical sections at 0.5µm was taken with 63x oil-immersion objective (NA 1.40; WD 0.19 mm, Zeiss).

### 2.3 Animals

Experiments were performed on WT and transgenic reporter mice on a C57BL6J background. Cre recombinase lines used included SST-IRES-Cre (Jackson Labs stock # 013044; Taniguchi et al., 2011), Pvalb-2A-Cre (Jackson Labs stock # 008069; (Hippenmeyer et al., 2005)), and VIP-IRES-Cre (Jackson Labs stock # 010908; (Taniguchi et al., 2011)). Homozygous Cre-expressing mice were mated with homozygous Ai3 (Jackson Labs Stock # 007903) or Ai14 (Jackson Labs Stock #007914) mice to create heterozygous transgenic mice with eYFP (YFP)- or tdTomato-labeled SST, PV, or VIP interneurons. Cells from at least three mice from each line were used to characterize FAP/YFPpost expression pattern.

### 2.4 Surgery

#### 2.4.1 Virus injection surgery

FAP/YFPpost virus (0.4 µL) or FingR virus (50nL; PSD95.FingR-GFP or Gephyrin.FingR-GFP)was stereotaxically injected into barrel cortex through a small craniotomy (bregma -0.9, lateral 3.00, depth 0.5 mm) in isoflurane-anaesthetized mice aged postnatal day (P11-15) using a Hamilton syringe (Hamilton; Reno, NV), Stoelting infusion pump 597 (Stoelting; Wood Dale, IL, Model #53210), and custom injection cannulas (Plastics One; 598 Phoenix, AZ). Mice were treated once with ketofen (5 mg/kg, Sigma-Aldrich; 599 St. Louis, MO), then allowed to recover in their home cage until weaning (P21),when they were moved to a new cage with their littermates.

#### 2.4.2 Window surgery

One week after virus injection, mice were isoflurane anesthetized and heads fixed using a custom-made nose clamp. Eyes were covered with ointment, hair was removed with Nair, and scalp was disinfected with povidone iodine. Scalp and periosteum was removed, and skull surface roughened by scraping with a slowly rotating dental drill. A thin layer of Krazyglue was applied to the skull before a custom-made head bracket was attached in the right hemisphere using Krazyglue and dental cement (Lang Dental, 1223PNK). The skull was carefully thinned around a 3mm diameter circle centered above the left hemisphere S1BF using a dental drill (Dentsply, 780044). After extensive thinning, the loose bone flap was detached using a microforcep. A glass window composed of 3mm diameter glass (Harvard Apparatus, 64-0720) attached to 5 mm diameter glass (Harvard Apparatus, 64-0700) was mounted above the exposed brain. The window was sealed with 3MTM VetbondTM. A chamber wall was built around the window with dental cement. Ketoprofen (3 mg/Kg) was given subcutaneously.

### 2.5 Tissue preparation

Eight to 15 days following virus injection, animals were anesthetized with isoflurane and transcardially perfused at mid-day using 20 mL phosphate buffered saline (PBS; pH 7.4) followed by 20 mL 4% paraformaldehyde in PBS (PFA; pH 7.4). Brains were removed, and postfixed overnight at 4°C in 4% PFA before transfer into 30% sucrose cryoprotectant. After osmotic equilibration brains were sectioned (50µm thick section) using a freezing-microtome. For tissue used in immunoelectron microscopy experiments, fixative contained EM-grade 4% PFA and 0.5% gluteraldehyde (Electron Microscopy Sciences, #15700 and #16000).

#### 2.5.1 Tissue preparation for confocal microscopy

Free-floating brain sections containing dTom-expressing cells were washed using PBS before 30 minute room temperature incubation with MG dye (300nM in PBS; (Pratt et al., 2017)). MG-dyed sections were then rinsed with PBS before mounting on glass microscope slides with Vectashield fluorescent mounting media (Vector Lab; Burlingame, CA). A subset of Ai3 x PV-Cre, SST-Cre, or VIP-Cre brain sections underwent GFP immunofluorescence staining to enhance YFP in presynaptic terminals before MG dye application. These sections were first blocked (10%NGS, 0.1% TritonX, 0.1M PBS), and incubated for 48 hour at 4°C with anti-chicken GFP primary antibody (1:2000 in blocking solution; Abcam AB13970; Cambridge, MA). Slices were rinsed with PBS then incubated with A488 anti-chicken secondary antibody (1:500, in blocking solution; Invitrogen A-11039; Carlsbad, CA). Similarly, subset of Ai3 x VIP-Cre brain sections underwent VAChT immunofluorescence staining to visualize cholinergic axon terminals using anti-VAChT primary antibody (1:2000 in blocking solution; Synaptic Systems 139 103; Goettingen, Germany) and A405 anti-rabbit secondary antibody (1:500; Invitrogen A-31556; Carlsbad, CA) before MG staining as described above.

#### 2.5.2 Tissue preparation for immunoelectron microscopy

Free-floating brain sections containing dTom-expressing cells were fixed in cold 2.0% PFA (Fisher, Pittsburgh, PA), 0.01 % glutaraldehyde (25% glutaraldehyde EM grade, Taab Chemical, Berks, England) in 0.01 M PBS (pH 7.3). The supernatant was removed and pelleted specimens were resuspended in soluble 3% gelatin in PBS (10-15µl; gelatin heated to 36°C prior to resuspension; Sigma, St. Louis, MO). Specimens were pelleted in this solution to concentrate them at the tip of the gelatin plug prior to gelling. Once solidified, gelatin pellets were fixed further in PFA/glutaraldehyde fixative. The pellets were infused with 20% polyvinylpyrrolidone 1.6 M sucrose buffered with 0.055 M sodium carbonate overnight (polyvinylpyrrolidone, sodium carbonate, Sigma, St. Louis, MO; sucrose, Fisher, Pittsburgh, PA), then frozen into small stubs in liquid nitrogen. Semi-thin (300 nm) sections were cut on a Leica Ultracut 7 with a Leica EM FC7 cryokit at -90° C (Leica Microsystems, Buffalo Grove, IL), stained with 0.5% Toluidine Blue in 1% sodium borate (Fisher, Pittsburgh, PA), and examined under a light microscope to identify a region of interest. Ultrathin sections (65 nm) were labeled with primary antibody (Chicken Anti-GFP, Abcam, #ab13970), and a colloidal gold labeled secondary (Donkey anti-chicken colloidal gold 6nm, Jackson Immunoresearch, #703-195-155). After rinsing in PBS, sections were fixed in 2.5% glutaraldehyde and rinsed in PBS and dH_2_O. The rinsing, sections were counterstained with 2% neutral uranyl acetate (Uranyl Acetate Dihydrate, Electron Microscopy Sciences, Hatfield, PA) and 4% uranyl acetate, coated with methyl cellulose.

### 2.6 Imaging

#### 2.6.1 Confocal microscopy

Sections were observed under a LSM 880 AxioObserver Microscope (Zeiss), using 63x oil-immersion objective lens (Plan-Apochromat, 1.40 Oil DIC M27) with the pinhole set at 1.0 Airy disk unit. Maximum image size was 1024 x1024 pixels. Zoom factor was set to 1, corresponding to a voxel dimension of 0.13µm x 0.13µm x 0.32µm in X, Y, and Z directions. Selected cell bodies were centered in the field of view (135µm x 135µm). Up to 100 images with a Z-interval of 0.32µm and 50% overlap between optical sections were acquired per stack. Fluorescence acquisition settings were as follows: YFP (ex514, em535, detection 517-553), dTom (ex561, em579, detection 561-641), MG/FAP (ex633, em668, detection 641-695). Optimal laser intensities for each channel were set for each cell independently, and images were collected to avoid pixel saturation (Supplemental Figure S1A,B). Well-isolated cells of interest were centered in the image frame and the Z-stack dimensions were set manually by tracking dTom labeled dendrites. Z-stacks typically ranged from 30-40µm for a given neuron.

##### 2.6.1.1 Cell Selection

FAP/YFPpost expression in L2/3 (~200-300µm below the pial surface) of the S1 barrelfield (S1BF) was confirmed by the presence of layer 4 barrels. Pyr cells were identified using morphological criteria, including the presence of a thick apical dendrite oriented toward the pial surface, pyramidal-shaped cell body, laterally projecting basal dendrites, a descending axon identified by its narrow diameter, and the ubiquitous presence of dendritic spines, particularly on higher-order branches. PV, SST, or VIP interneurons were identified in tissue collected from Ai3 x PV-Cre, SST-Cre, or VIP-Cre double transgenic lines using YFP expression. Both PV and SST neurons sometimes showed dendritic spines, although these were at a substantially lower density than in Pyr neurons. Only cells expressing dTom and punctate FAP/YFP signal were selected for quantitation. dTom-expressing neurons that did not exhibit membrane-localized fluorescence, or showed diffuse and non-punctate signal were excluded from analysis. In almost all cases, selected cells included the entire soma in the image dataset. Because cortical dendrites are > 200µm long and could lie outside the imaged area, only a fraction of the dendritic arbor was collected and analyzed.

#### 2.6.2 Electron microscopy (EM)

Sections were examined on a JEOL 1400 transmission electron microscope (JEOL Peabody, MA) with a side mount AMT 2K digital camera (Advanced Microscopy Techniques, Danvers, MA).

#### 2.6.3 Two-photon imaging

Mice were anesthetized with 1.5 % isoflurane and mounted under a Femtonics FEMTO2D microscope. Layer 1/2 dendrites expressing dTom and YFPpost were visualized under a 63x objective using 950 nm excitation (Spectra-Physics Mai Tai HP; Santa Clara, CA) with simultaneous detection of dTomato and YFPpost using red and green PMTs, respectively. Single-plane 60×60µm (1000×1000 pixel) linescan (15x averaging) images were acquired using MES software (Femtonics, v.5.2878). The raw intensity matrix for each channel was converted to a grayscale image in MATLAB (MathWorks, R2017a). Channels were overlaid and brightness/contrast adjusted using Photoshop 6.0 (Adobe). Because specimens were fixed and sectioned (and thus truncated), very distal segments of a given neuron’s dendritic arbor were not always captured during imaging.

### 2.7 Image analysis

#### 2.7.1 Imaris segmentation

Carl Zeiss image files were imported into Imaris version 8.4 equipped with the Filament Tracer plugin (Bitplane; Zurich, Switzerland). Typically, the dTom cell fill was used to create a 3D cell-surface rendering using a combination of surface and filament objects. For a subset of FAPpost-expressing VIP and SST cells, YFP fluorescence allowed for better detection of the dendritic arbor than dTom. For cell and neurite reconstruction, FAP/YFPpost puncta were first reconstructed as 3D structures using “surface objects” (to outline puncta borders). To ensure that all puncta were detected, fluorescent signal was enhanced to enable visualization of both bright and weak puncta (Figure S1C). Subtraction of background fluorescence was performed by masking cell-associated fluorescence (Figure S1D). Due to imaging limitations, only puncta larger than 3 voxels (~0.024µm3) were counted, potentially undercounting very small synapses below this detection threshold. Large puncta that potentially reflected smaller, fused synapses were separated into multiple objects with an estimated 0.5µm diameter using the “split touching objects” function (Supplemental Figure S1E). Thus, large puncta were potentially separated into multiple smaller synapses, a process that could increase the absolute number of detected synapses. Indeed, it is unclear for larger synapses whether these should be counted as a single synapse with multiple active sites and post-synaptic specializations (Tang et al., 2016), or combined into one large synapse (such as the giant synapses observed at the Calyx of Held). Using the dTom signal to identify cell surface boundaries, puncta were digitally associated with the plasma membrane if their edges lay within 0.5µm from the cell surface (<1µm for spiny dendritic regions >30µm from the soma center). Puncta 0.5µm below cell surface were attributed as cytosolic puncta and not included for analysis. Puncta “objects” were then converted into puncta “spots” (with automatic intensity max spot detection thresholds and a 0.5µm estimated-diameter) using “object” centroids in Imaris.

#### 2.7.2 Puncta quantification

Puncta were quantified for individual dendritic branches and separately averaged for different branch orders. Pyr neurons had only one apical branch segment. The number and length of 2° and higher-order branches analyzed could vary across cells, depending upon cell anatomy and image acquisition. For single-cell dendritic puncta density averages (Figure 4,5), values for the Pyr 1° apical dendrite were not included, because this compartment showed a significantly higher density and appeared to be a unique compartment of the neuron that was contiguous with the soma. Although the axon is easily distinguished in Pyr neurons based upon its basal location and small diameter, interneuron axons can be hard to distinguish. Thus, Pyr axons were excluded from analysis of dendritic puncta densities, but may have been included for some interneurons, particularly for VIP neurons where the axon exits from the basal dendrite distant from the soma (Pronneke et al., 2015).

**Figure 4:**
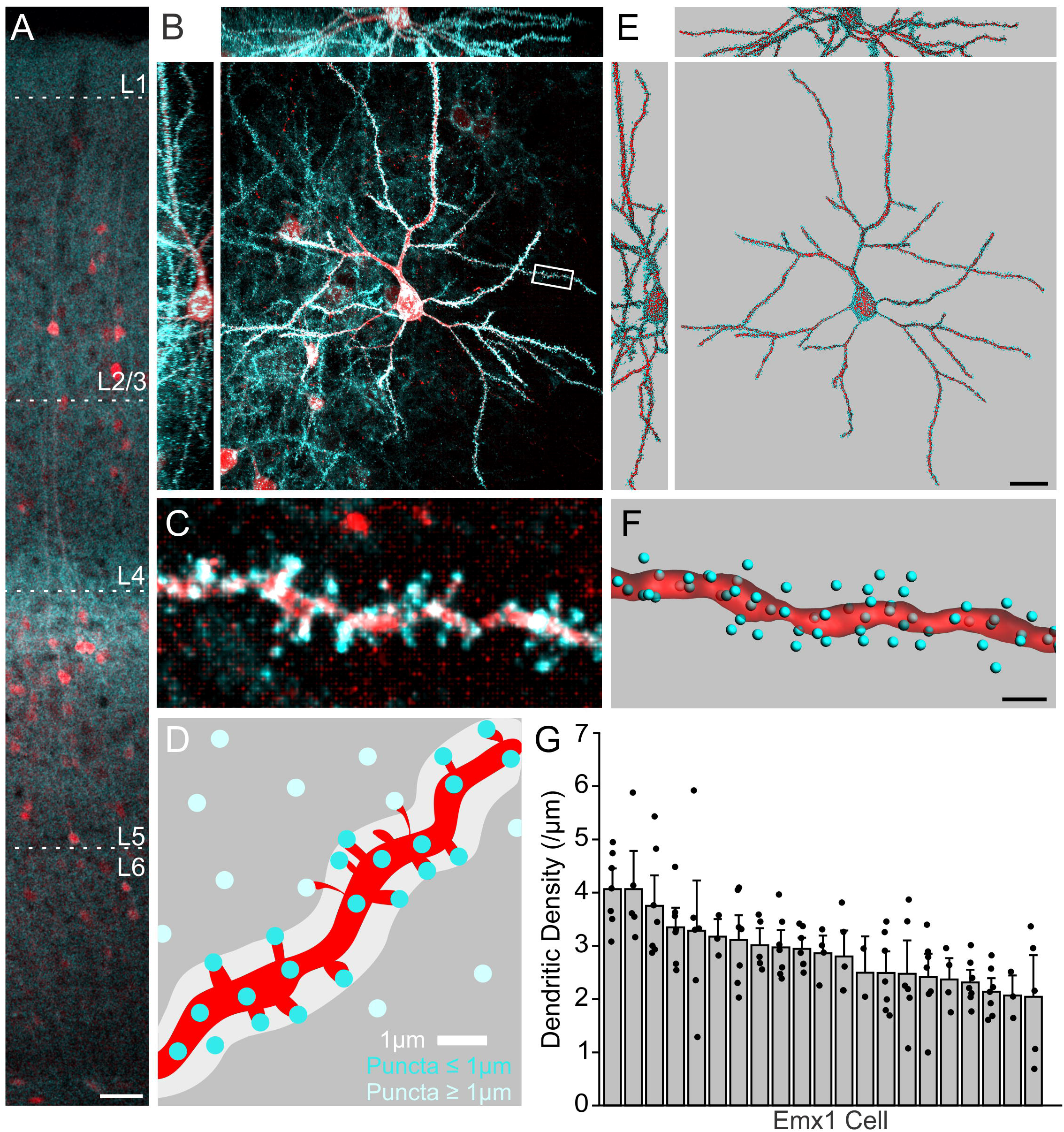
YFPpost fluorescent puncta quantitation in L2/3 Pyr cell dendrites. (A) YFPpost expression across the cortical column. Scale=50µm. (B) L2/3 Pyr expressing YFPpost. (C) YFPpost fluorescent puncta on a dendritic shaft and spines (zoom from box in *B*). (D) Schematic for dendritic puncta assignment (blue, assigned puncta <1.0µm from shaft surface; light blue, unassigned puncta). (E) 3D-rendering of Pyr neuron (red) with assigned puncta. Scale=20µm. (F) Dendrite from *C* with assigned puncta. Scale=2µm. (G) Mean YFPpost puncta density for individual neurons (grey bars, +SEM) averaged from individual apical and basal dendritic branch densities (black dots). n=21, N=4. See also Supplemental Figure S1 and S2.

**Figure 5:**
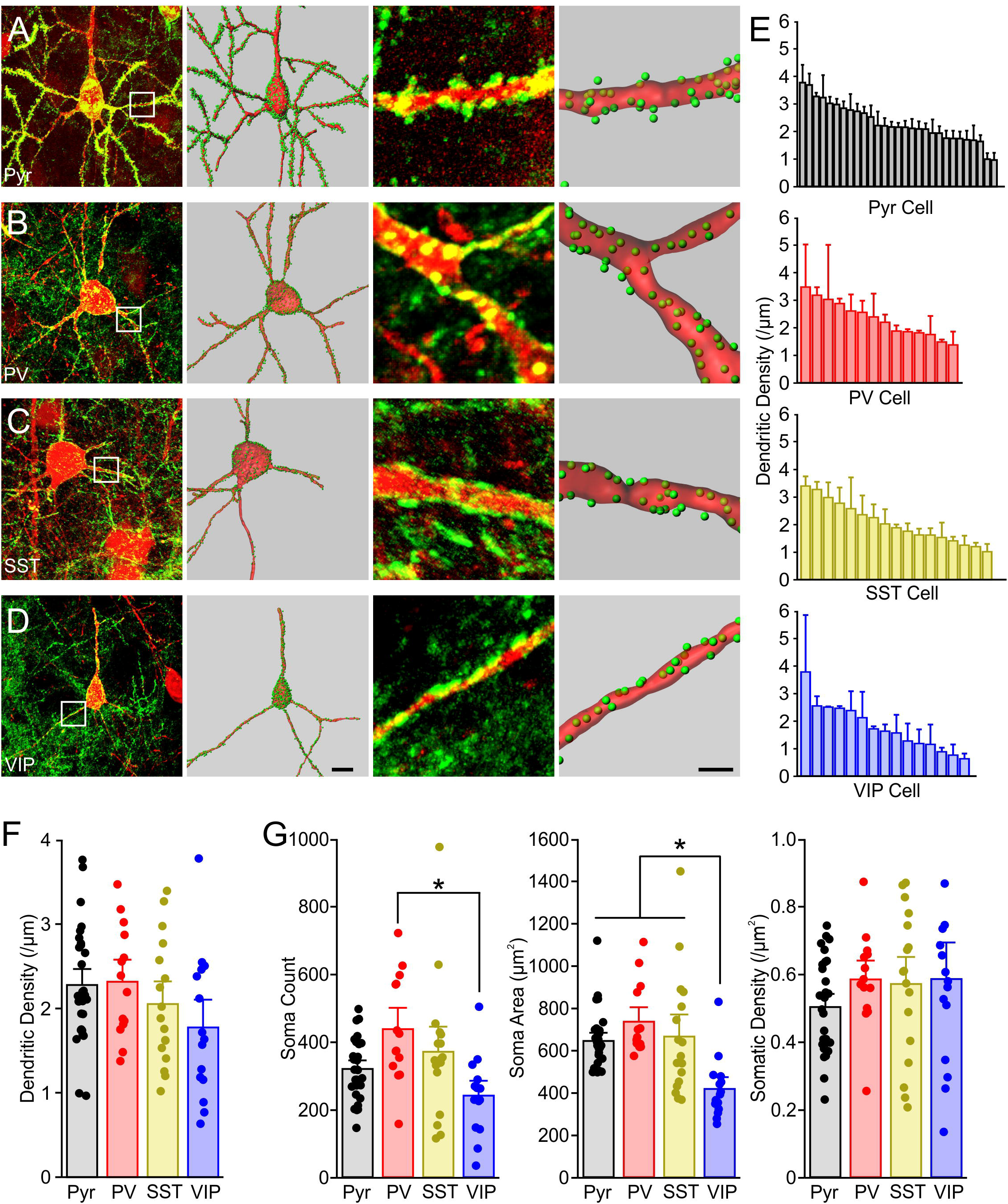
Cell-type specific synapse quantitation for L2/3 Pyr, PV, SST, and VIP neurons. (A) From left: Confocal stack of example FAPpost Pyr neuron; 3D-rendering of the neuron and puncta assignment; zoom of dendritic segment; 3D-rendering of segment. Scale=10µm, 2µm. (B,C,D) As in *A* but for PV, SST, and VIP neurons. (E) Dendritic puncta density across individual Pyr, PV, SST, and VIP cells. Bars are cell means (+SEM). (F) Dendritic puncta density does not differ across cell types. Dots are individual cell means and bars are all-cell means+SEM. (ANOVACellType: F(3,71)=1.7, *p*=0.1). (G) Soma size differs but puncta density is constant across cell types. Left: total somatic puncta are significantly greater in PV neurons (ANOVACount: F(3,71)=5.2, *p*=0.003). Middle: VIP neurons are smaller than other interneuron types (ANOVASurfaceArea: F(3,71)=8.0, *p*<0.001). Right: Somatic puncta density is conserved across cell types (ANOVADensity: F(3,71)=0.96, *p*=0.4). *Posthoc Tukey multiple comparison test, *p*<0.05. See Supplemental Figure S3, S4, and Supplemental Table S1 for n’s.

#### 2.7.3 Input assignment

Puncta spots were assigned to a specific presynaptic input using a distance threshold of 0.15µm from spot centroid to presynaptic neurite 3D-reconstructions. Presynaptic neurite reconstructions were created using automatic background subtraction thresholding of the presynaptic (PV, SST, or VIP) YFP channel, a split-touching objects diameter threshold of 1µm, and 5 voxel minimum area settings in Imaris. For confocal stacks where presynaptic neurite YFP signal exhibited z-axis related signal drop-off, neurite reconstructions using automatic settings were generated separately for superficial (first ~20µm) and deeper optical sections of the stack. In such cases, both sets of presynaptic neurite reconstructions were visually examined for comparable density and size profiles.

Methods using pre- and postsynaptic neurite colocalization may be confounded by false positives (where neurites are near the soma but do not synapse onto it) and false negatives (where a given neurite is associated with two or more postsynaptic sites that are conflated into a single crossing point). Indeed, in many cases, neurites made extended contacts with the soma surface that might include a single or multiple postsynaptic sites. Using postsynaptic puncta to differentiate multiple synapses along an extended region of presynaptic neurite puncta enabled a more accurate estimate for true synapse number (Supplemental Figure S5).

Because distance parameters used to identify convergent signals could be digitally adjusted, we explored this space to establish a maximum distance for input detection of 0.15µm. This was below the diffraction limit for our confocal images. As expected, use of larger distance thresholds for detecting inputs resulted in a substantial increase in the number of assigned puncta. These values were out of range for other published values for inhibitory synapse density and provided confirmation that smaller distance thresholds were more stringent and likely to be more accurate.

For SST-input assignment onto SST cells, the same maximum distance of 0.15µm was used. Since the postsynaptic cell also expressed YFP, all neurites within 0.1µm of the analyzed postsynaptic cell were excluded as potential inputs because they could not be differentiated from the parent dendrite.

### 2.8 Electrophysiology

FAPpost-injected mice were sacrificed at age P20-25 by brief isoflurane anesthesia and decapitation. Coronal slices (350µm thick) were prepared in regular ice-cold artificial cerebrospinal fluid (ACSF) composed of (in mM): 119 NaCl, 3.5 KCl, 1 NaH_2_PO_4_, 26.2 NaHCO_3_, 11 glucose, 1.3 613 MgSO_4_, and 2.5 CaCl_2_ equilibrated with 95%/5% O_2_/CO_2_. Slices recovered in the dark at room temperature for 60 minutes before transfer to an electrophysiology rig where they were perfused with ACSF containing 1µM tetrodotoxin (Tocris, UK) to silence spontaneous activity. The injection site was identified by dTom fluorescent cell bodies using an Olympus light microscope (BX51WI). Pyr-targeted recordings (4-5 animals per group) were done in the absence of MG dye, since we were interested in whether *in vivo* expression of the FAPpost construct would influence synaptic function and MG was never applied before tissue fixation for anatomical analysis. Borosilicate glass electrode resistance was 4-8 MΩ. Electrode internal solution was composed of (in mM): 130 cesium gluconate, 10 HEPES, 0.5 EGTA, 8 NaCl, 10 Tetraethylammonium chloride (TEA-Cl), 4 Mg-ATP and 0.4 Na-GTP, pH 7.25-7.30, 280-290 mOsm. Trace amounts of AlexaFluor488 were included in the internal solution to confirm that targeted cells had pyramidal-like morphologies. Electrophysiological data were acquired using a Multiclamp 700B amplifier (Axon Instruments; Foster City, CA) and a National Instruments acquisition interface (National Instruments; Austin, TX). The data were filtered at 3 kHz, digitized at 10 kHz and collected by Igor Pro 6.0 (Wavemetrics; Lake Oswego, Oregon). After forming a GΩ seal, negative pressure was applied to the cell to enter whole-cell mode, and following 2-3 minutes acclimation time, miniature excitatory postsynaptic currents (mEPSCs) were collected at -70 mV holding potential for 5 minutes. Holding potential was slowly raised to 0 mV over an additional minute, and following 1 minute acclimation time, miniature inhibitory postsynaptic currents (mIPSCs) were then collected. Traces were analyzed using MiniAnalysis (Synaptosoft Inc., NJ), with a 7pA minimal amplitude cut-off. One hundred randomly selected events for each cell (Pyr dTom- and dTom+) were used to create cumulative probability histogram.

### 2.9 Statistical Analysis

All reported values are mean ± SEM, unless otherwise stated. Dendritic puncta density is mean spot density per linear dendritic segment for a given cell. Soma density is total somatic spot count divided by soma surface area. Density distributions were tested for normality both within and across cells using the Shapiro-Wilk normality test. Within cells, all but two Pyr cells had normally distributed dendritic puncta densities. For these two cells, median dendritic puncta density was used to represent these cell’s average dendritic puncta densities. Mean dendritic puncta density was used for all other cells. One-Way ANOVA was used to detect cell-type group differences. One-Way repeated measures (RM) ANOVA was used to detect dendritic segment-level dependent puncta density within cell-type (p < 0.05). Two-Way ANOVA was used to detect effects of animal, age, or days post infection (DPI) across cell-types. Pearson’s correlation was employed to test the relationship between Pyr soma surface area and synapse density. Two-Way RM ANOVA was used to detect differences in the proportion of input-assigned synapses across Pyr compartments and input-types (p < 0.05). Post-hoc Tukey’s multiple comparison testing was performed to identify significant group mean differences for anatomical data. For physiological data, unpaired Student’s T-test was used to identify significant differences in mean mEPSC and mIPSC amplitude and frequency, and Kolmogorov-Smirnov test was used to test for differences in amplitude distributions (p < 0.05). All analyses were performed using Origin 2017 statistical software (OriginLab, Northampton, MA).

## 3 Results

### 3.1 Fluorophore targeting to postsynaptic sites

Neuroligins are ubiquitously expressed at postsynaptic sites (Bemben et al., 2015). We took advantage of the pan-synaptic localization of a NL-1 based tether (Kim et al., 2011) to direct an extracellular fluorophore to postsynaptic sites for individual neurons in the intact cortex. Because the trans-synaptic protein-protein interactions involved in neuroligin-neurexin complexing and GRASP have been linked to synaptogenesis and stabilization (Scheiffele et al., 2000; Tsetsenis et al., 2014), we employed an intact fluorophore for postsynaptic labeling.

We constructed NL-1-based synaptic targeting constructs where the extracellular portion of NL-1 was replaced with either FAP or YFP, under the control of the human synapsin promoter (Figure 1A-C). Constructs included a cell-filling dTom and were engineered so that they could be activated by Cre-dependent recombination or could be expressed ubiquitously in all neurons. AAV-packaged viral constructs were injected into mouse primary somatosensory (barrel) cortex (Figure 1D), and transduced cells were identified in fixed tissue specimens by both dTom and FAP or YFP expression without signal amplification. Confocal imaging revealed punctate FAP or YFP signal on the cell body and dendrites of L2 Pyr neurons (Figure 1E-L; Supplemental Movie S1). Puncta were observed on both dendritic shafts and spines, where ~90% of dTom-filled spines exhibited FAP/YFPpost puncta. Some puncta could be detected in close proximity to a well-isolated dendrite that lacked an apparent spine (Figure 1L asterisk), suggesting that not all spines were visible in fluorescence microscopy images.

**Figure 1:**
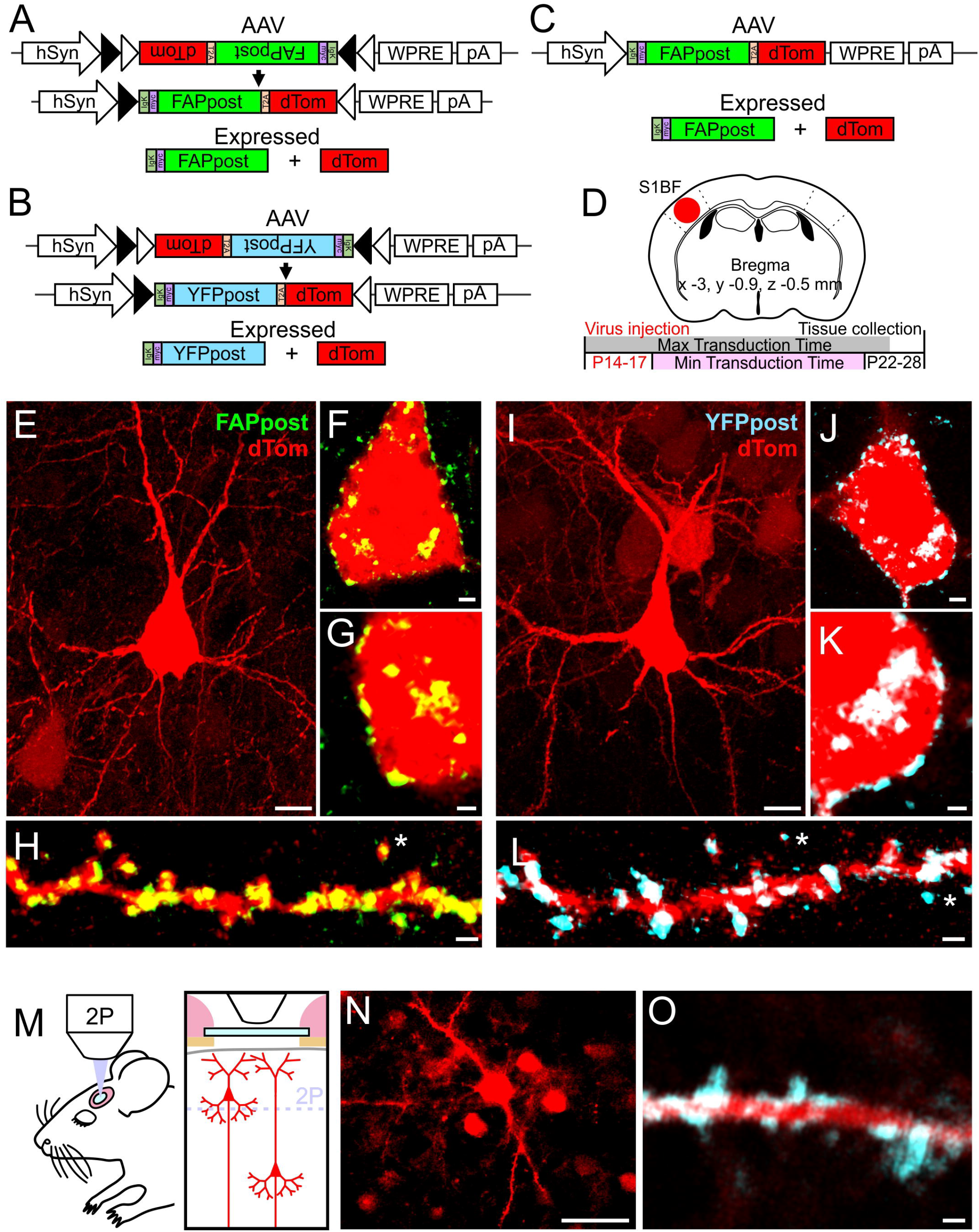
Construct design and expression in mouse somatosensory (S1 barrel fields) cortex. (A) Fl-FAPpost construct design. Human synapsin promotor (hSyn) driving Cre-dependent FAPpost and dTom, separated by a 2A sequence for independent localization. (B) Fl-YFPpost construct. Cre-dependent YFPpost and dTom expression. (C) FAPpost construct. Cre-independent FAPpost and dTom expression. (D) Virus injection coordinates into WT or Emx1-Cre mouse and experimental time-course. (E) Confocal stack of L2 pyramidal cell transfected with FAPpost. Scale=10µm. (F) Optical section of FAPpost puncta on soma of cell in *E*. Scale=2µm. (G) Zoom of *F*. Scale=1µm. (H) FAPpost labeled spiny dendrites. Scale=1µm. (I) Confocal stack of L2 pyramidal cell transfected with YFPpost. (J) Optical section of YFPpost puncta on soma of cell in *I*. Scale as in *F*. (K) Zoom of *J*. Scale as in *G*. (L) YFPpost labeled spiny dendrites. Scale=1µm. (M) *In vivo* 2P imaging schematic. (N) Single-plane 2P image of L2 Pyr cells transfected with YFPpost. Scale=30µm. (O) Single-plane 2P image of YFPpost labeled spiny dendrite in L1. Scale as in L+H. See also Supplemental Movie S1.

*In vivo* 2-photon (2P) imaging of YFPpost-transfected dendrites in mouse S1 revealed that punctate YFP signal was associated with both dendritic shafts and spines (Figure 1M-O). Because FAPpost fluorescence required the addition of MG fluorogen, and *in vivo* imaging was carried out using a cranial window with a glass coverslip after several days of recovery, it was not possible to image FAPpost expression *in vivo*. In addition, the excitation and detection parameters for FAPpost are not optimized for 2P imaging using a conventional laser set-up. However, our results indicate that YFPpost is bright enough for detection of puncta in living tissue. Overall, we find that NL-1 tethered fluorophores can be detected in both fixed and living brain tissue without signal amplification, with punctate expression that localizes to sites of synaptic input.

### FAPpost localizes to both excitatory and inhibitory synapses

The distribution of puncta at somatic, shaft, and spine locations in L2 Pyr neurons was consistent with labeling of both excitatory and inhibitory synapses. To verify this, we carried out immunocytochemistry for PSD-95, a postsynaptic scaffolding protein that anchors glutamate receptors, and for gephyrin, a postsynaptic scaffolding protein that complexes with GABAa receptors. FAPpost-expressing cultured neocortical neurons showed colocalization of both PSD-95 and gephyrin with FAPpost puncta (Figure 2A-D,F-I), suggesting that these constructs can target both excitatory and inhibitory synapses.

To further confirm synaptic localization, we used immuno-EM against YFP from YFPpost expressing neurons in S1. Gold particles were concentrated at the synapse for asymmetric and symmetric synapses (Figure 2E,J), consistent with YFPpost localization at excitatory and inhibitory synapses. A comparison of cross-sectional synapse diameters showed that YFPpost expression did not alter synapse size (250±105nm) compared to unlabeled synapses (241±112nm).

**Figure 2:**
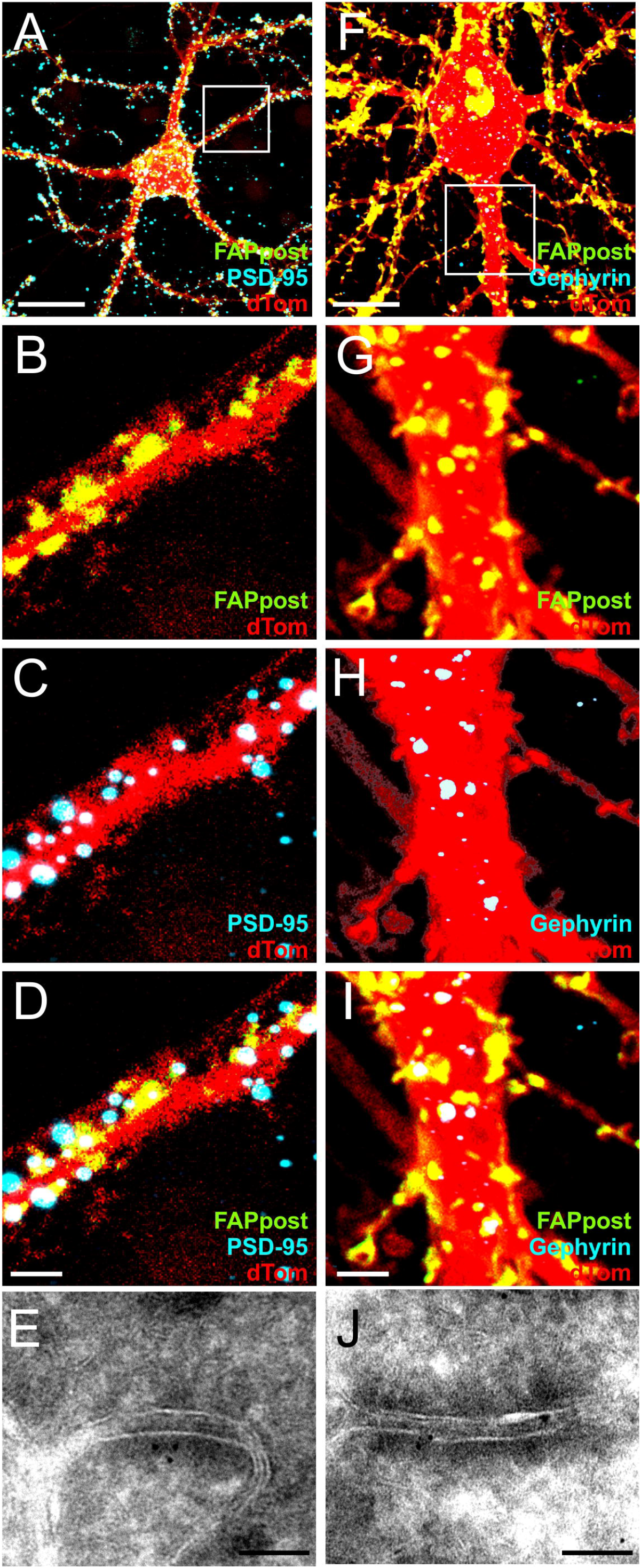
FAPpost colocalizes with excitatory and inhibitory synaptic markers. (A) Cultured neocortical neurons transfected with FAPpost and immunostained for PSD-95. (B) FAPpost labeled dendrite from boxed area in *A*. (C) Dendrite and PSD-95. (D) Colocalization of FAPpost and PSD-95 puncta. (E) GFP-immunogold labeling of YFPpost at an asymmetric synapse in EMX1-Cre mouse cortical tissue visualized using TEM. (F) As in *A,* but for a different transfection and immunostained for gephyrin. (G) FAPpost labeled dendrite from boxed area in *F*. (H) Dendrite and gephyrin. (I) Colocalization of FAPpost and gephyrin. (J) GFP-immunogold labeling of YFPpost at symmetric synapse in EMX1-Cre mouse cortical tissue visualized using TEM. Scale *A+F*= 10µm, *B-D* and *G-I* = 2µm, *E+J*=100nm.

The association of FAP/YFPpost with both inhibitory and excitatory synaptic markers indicates that these reagents can label multiple synapse types, and that they can be a useful marker for broad-scale synaptic quantitation across an entire neuron.

### 3.3 FAPpost does not disrupt synaptic function

Overexpression of other genetically-encoded synaptic proteins has been associated with elevated synapse density and abnormal electrophysiological properties. For example, increased anatomical synapse density and mEPSC frequency has been observed with overexpression of GFP-tagged PSD-95, gephyrin, or intact NL-1 (Chubykin et al., 2005; El-Husseini et al., 2000; Gross et al., 2013; Prange et al., 2004). Trans-synaptic interactions for split protein indicators have been shown to increase binding affinities for the tagged proteins, and irreversible GFP-reconstitution can perturb synapse stability and organization (Tsetsenis et al., 2014; Yamagata and Sanes, 2012).

Electrophysiological recordings can be a sensitive way to survey alterations in synaptic function, independent of anatomical quantitation from fluorescence images. To test whether FAPpost expression was associated with altered mEPSC and mIPSC properties, adjacent untransfected and dTom+, FAPpost transfected cells were targeted for whole-cell recordings (Figure 3).

**Figure 3:**
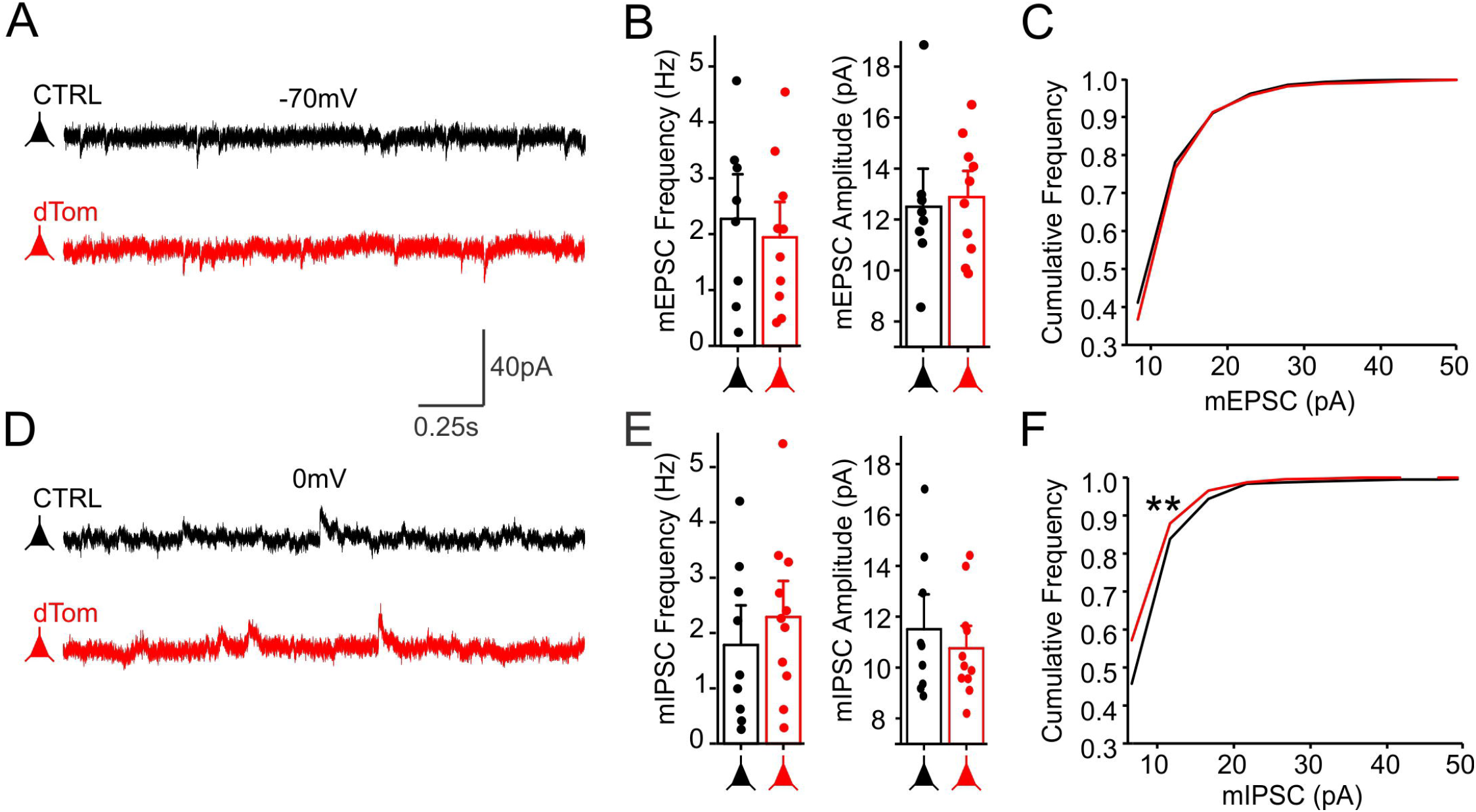
FAPpost synaptic localization does not alter mEPSC and mIPSC properties. (A) Example voltage-clamp traces from an untransfected (black) and neighboring FAPpost-expressing (red) L2/3 Pyr cell showing mEPSCs. (B) Comparison of mean mEPSC frequency (ANOVAFrequency: F(1,16)=0.2, *p*=0.6) and amplitude (ANOVAAmplitude: F(1,16)=0.1, *p*=0.8) indicate no difference. (C) Cumulative distribution histogram of mEPSC amplitudes (Kolmogorov-Smirnov Test, D=0.05, *p*=0.16). (D) Example voltage-clamp traces from an untransfected (black) and neighboring FAPpost-expressing (red) L2/3 Pyr cell showing mIPSCs. (E) Mean mIPSCs frequency (left) of untransfected and dTom cells were not significantly different (ANOVAFrequency: F(1,18)=0.6, *p*=0.4). Mean mIPSC amplitude (right) of untransfected and dTom cells were not significantly different (ANOVAAmplitude: F(1,18)=0.5, *p*=0.5). (F) Cumulative distribution histogram of mIPSC amplitudes shows a small but significant shift in dTom mIPSC amplitudes (Kolmogorov-Smirnov Test, D=0.15, *p*<0.0001). n=8-11, N=7.

FAPpost expression did not alter mean mEPSC frequency or amplitude (Figure 3B; frequency untransfected 2.3±0.5 Hz vs FAPpost 1.9±0.4 Hz; amplitude untransfected 12.5±1.0 pA versus FAPpost 12.8±0.7 pA). Furthermore, mean mIPSC frequency and amplitude were not significantly different (Figure 3E; frequency untransfected 1.8±0.5 Hz versus FAPpost 2.3±0.4 Hz; amplitude untransfected 11.5±0.9 pA vs FAPpost 10.8±0.6 pA), although a small reduction in the frequency distribution of mIPSCs was observed (Figure 3F). Thus, expression of postsynaptic fluorophores using NL-1 targeting sequences can be a non-invasive way to identify and quantitate synaptic distributions without altering synaptic function.

### 3.4 High-throughput synapse quantitation

To facilitate quantitative fluorescence analysis of synapses in neurons from brain tissue, we applied an efficient and scalable analysis pipeline with automated synapse detection and assignment. Sparse viral transduction of YFPpost in Pyr neurons from primary somatosensory (barrel) cortex of Emx1-Cre transgenic mice revealed well-isolated Pyr neurons decorated with bright, YFP puncta across the cell surface (Figure 4A-C). Using Imaris image analysis software, neural surfaces were rendered and puncta assigned to an individual neuron for quantitative analysis (Figure 4D-G). Because dendritic spines were not always visible from the dTom fill, YFPpost puncta that were 1µm from the parent neurite were digitally assigned to a given cell for density calculations. This semi-automated approach enables high-throughput and cell-type specific synapse identification and quantitative analysis (Supplemental Figure S1).

Reconstructions of fixed specimens yielded 200-950µm of continuous dendritic segment for analysis. We observed substantial heterogeneity in puncta density across individual L2 Pyr neurons, with two-fold variance across cells (mean range, 2.0 to 4.0 puncta/µm), and nearly 10-fold variance (0.7 to 5.9 puncta/µm) across dendritic branches (Figure 4G). We found no relationship between mean puncta density and total dendritic length for a given neuron, suggesting we captured the true variability of puncta density across L2 Pyr neurons. Overall, YFPpost puncta densities across L2 Pyr dendrites (2.9+0.6 puncta/µm) were similar to previous estimates of synapse density (Gulyas et al., 1999; Hersch and White, 1981; Holtmaat et al., 2005; Kasthuri et al., 2015; Villa et al., 2016a), providing validation of this high-throughput analytical approach. In addition, a direct comparison of puncta densities using FAP/YFPpost constructs in L2/3 PV-Cre expressing neurons showed that these reagents yield quantitatively similar results, indicating that the constructs are interchangeable (Supplemental Figure S2).

### 3.5 Cell-type specific characterization of synapse density

It is unknown whether synapse density systematically differs according to cell type, a question that can be addressed by quantitative analysis. In addition, establishing the normal range of synapse densities across different cell types can provide an important benchmark for assessing learning or disease-related changes. We thus evaluated the distribution of putative synapses across four different neocortical neuron types in mouse barrel cortex: L2/3 Pyr, PV, SST, and VIP neurons (Figure 5A-D). FAPpost-expressing neurons were identified using morphological parameters and overlap with Cre-dependent YFP expression in Ai3 reporter lines.

Overall, dendritic puncta densities were remarkably similar across different cell types (Figure 5E-F; PV: 2.3±0.2/µm, SST: 2.1±0.2/µm, VIP: 1.8±0.2/µm, and Pyr: 2.3±0.1/µm excluding the 1° apical dendrite; see Methods). PV neurons exhibited the highest somatic puncta counts (Figure 5G), but when corrected for soma surface area, all cell types showed similar puncta densities. Although there are multiple distinct subclasses within the PV, SST, and VIP group (see for example: (Jiang et al., 2015; Kubota et al., 2016; Munoz et al., 2017; Pronneke et al., 2015) we did not observe puncta density-defined subgrouping of neurons. Rather, puncta densities for individual cells within a class were distributed along a continuum (Figure 5E).

Variability in puncta density for any cell type could not generally be explained by cell-type, age, days post-infection, or animal-to-animal differences as neurons with a range of puncta densities could be found in the same animal (Supplemental Figure S3 and Supplemental Table S1). VIP and Pyr neurons displayed the largest within-group variability (~4-fold, Figure 5F), and Pyr neurons showed a modest influence of animal or litter on puncta density (Supplemental Figure S3; see also (Chandrasekaran et al., 2015)). This is consistent with anatomical and electrophysiological response variability that has been described for these groups of neurons in L2/3 (Pronneke et al., 2015; Tasic et al., 2016; Tyler et al., 2015; van Aerde and Feldmeyer, 2015; Yamashita et al., 2013; Yassin et al., 2010), and may reflect both developmental and molecular heterogeneity.

Prior reagents have attempted to leverage fluorescence imaging for comprehensive synapse labeling, including mGRASP, PSD95-or Gephyrin-FingRs (Gross et al., 2013; Kim et al., 2011). To compare the distribution of YFPpost synapses with a separate synapse labeling method, we quantitatively examined synapse labeling using FingR-GFP PSD95 and gephyrin intrabodies and compared them to YFPpost results for a molecularly defined cell type, L2 SST cells.

We observed almost 20-fold variability in mean puncta density across labeled dendritic segments for both PSD95 or gephyrin FingRs in SST neurons, a phenomenon that was visible even when comparing primary dendritic branches (Supplemental Figure 4). The wide range of synapse densities was most influenced by neurons with very low numbers of labeled synapses, despite nuclear labeling indicating FingR expression. Similar, though slightly less variability (10-fold) was present for YFPpost-labeled SST cells from the same tissue. Compared to FingRs, YFPpost expression showed more faint and diffuse membrane fluorescence, although well-isolated puncta could be easily detected.

Overall, the sum of the mean FingR PSD95- and gephyrin labeled-puncta (mean density of 1.12±0.47 synapses/µm) was similar to the total puncta numbers labeled with YFPpost (mean density 1.45±0.85 synapses/µm) for the SST neurons in this dataset. We note that FAP/YFPpost puncta could be detected at other synapse types, for example cholinergic synapses (Supplemental Figure S4). Because FAP/YFPpost constructs may label multiple synapse types and because some bonafide excitatory and inhibitory synapses do not contain gephyrin or PSD95 (Cane et al., 2014; Sassoe-Pognetto et al., 1994; Villa et al., 2016b; Viltono et al., 2008), we do not expect that the total synapse number calculated from YFPpost detection will be the simple sum of gephyrin- and PSD95-labeled synapses.

### 3.6 Compartment-specific analysis of synapse density in Pyr cells

Comprehensive labeling of synapses enables analysis of synapse organization across somatic, dendritic, and axonal compartments. We took advantage of the polarity of Pyr neurons to determine whether synapses were uniformly distributed across their somatic, apical and basal compartments.

L2 Pyr soma surface area varied more than 2-fold, consistent with prior observations that neocortical pyramidal neurons can have different mean soma sizes (van Aerde and Feldmeyer, 2015) associated with distinct developmental origins and projection targets (Deitcher et al., 2017; Tyler et al., 2015; Yamashita et al., 2013; Zaitsev et al., 2012). Soma puncta density was not correlated with soma size, although higher somatic and dendritic puncta densities were positively correlated (Supplemental Figure S3H-J).

Apical and basal dendrites of cortical Pyr neurons are distinct processing structures (Larkum et al., 2009; Larkum et al., 2007; Spruston, 2008) and may be targeted by different presynaptic inputs. We thus compared puncta distribution across primary (1°) through quaternary (4°) branches of apical and basal compartments within ~200µm of the soma (Figure 6). Overall, puncta density for higher-order apical and basal dendrites of pyramidal neurons was similar.

**Figure 6:**
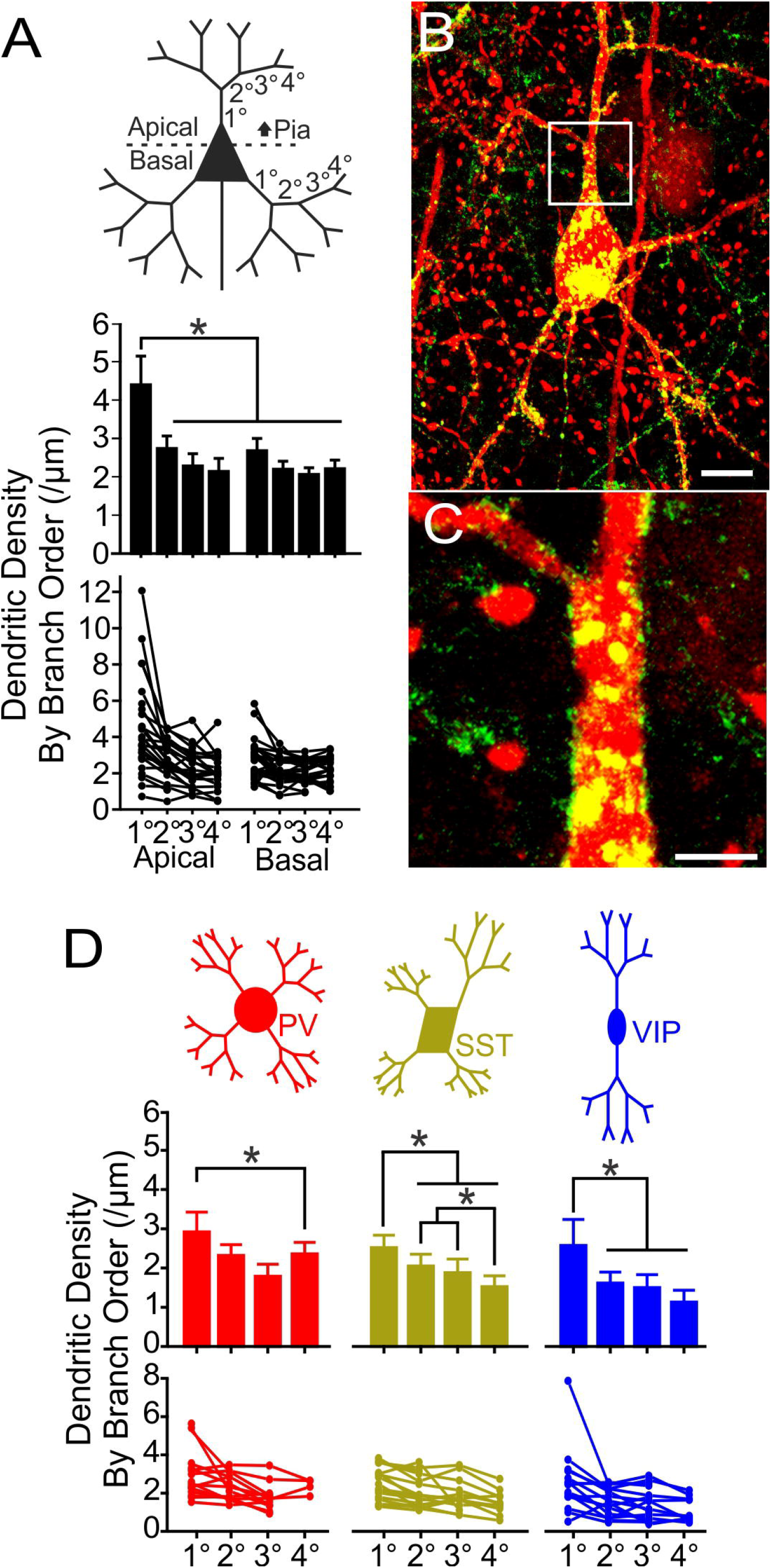
The primary apical dendrite of Pyr neurons has a high synapse density. (A) (top) Schematic of Pyr branch orders analyzed. (middle) Mean FAPpost synapse density across 1°-4° Pyr branches (bar +SEM). (bottom) Individual cell values plotted as connected lines. Repeated measures (RM) ANOVAPyr: F(7,112)=17, *p*<0.0001. (B) Confocal stack of example Pyr cell. Scale=10µm. (C) Zoom of boxed region in *B*. Scale=3µm. (D) As in *A*but for comparison of PV (red), SST (yellow), and VIP (blue) synapse densities by branch. RM ANOVAPV: F(3,12)=4.7, *p*=0.02. RM ANOVASST: F(3,30)=21, *p*<0.0001. RM ANOVAVIP: F(3,27) =6.2, *p*=0.002. All data shown in graphs (Pyr n=29, N=12; PV n=14, N=3; SST n=17, N=5; VIP n=15, N=3). Statistical comparisons performed on balanced data where all branch orders were present and could be tested (Pyr n=17, N=8; PV n=5, N=2; SST n=12, N=5, VIP n=10, N=3). *Tukey post-hoc pairwise comparison test, *p*<0.05.

Puncta density in the 1° apical dendrite was significantly elevated compared to other dendrites (Figure 6A-C). The 1° apical dendrite showed a mean of 4.4±0.5 puncta/µm, greater than all other apical branches (2°=2.7±0.2/µm; 3°=2.3±0.2/µm; 4°=2.2±0.2/µm) and basal branches (1°=2.6±0.2/µm; 2°=2.2±0.1/µm; 3°=2.1±0.1/µm; 4°=2.2±0.1/µm). The smooth, aspiny region of the apical dendrite has not been well-studied in prior studies, likely due to the use of spines as a proxy for synapses.

Inhibitory neurons are frequently multipolar, and a clear distinction between the apical dendrite and other dendrites was not always apparent. Analysis of PV, SST, and VIP neurons indicated their 1° dendrites also exhibited a higher density of puncta than higher-order dendritic branches (Figure 6D), although this difference was less pronounced than that observed for the Pyr 1° apical dendrite.

### 3.7 Identifying presynaptic cell identity for labeled synapses

Synaptic location along the dendrite and presynaptic input identity are primary determinants of how synapses can alter neural function. The far-red emission of MG-binding FAPs is particularly useful for determining input identity, since it can be combined with other commonly-used fluorophores. Cre-driver lines were used for the selective and comprehensive fluorescence labeling of discrete classes of neocortical inhibitory neurons (Hippenmeyer et al., 2005; Taniguchi et al., 2011). Initially we focused on YFP-labeled PV, SST, and VIP inputs to Pyr soma, since data from prior EM and light-microscopy analyses could corroborate our results (Di Cristo et al., 2004; Hill et al., 2012; Kubota et al., 2016; Kubota et al., 2015; Melchitzky and Lewis, 2008; Tamas et al., 2000; Zhou et al., 2017).

The smooth postsynaptic membrane of Pyr soma enabled us to evaluate different distance thresholds for aligned PV or SST neurites with post-synaptic FAPpost puncta. First, using only neurite-associations with the postsynaptic soma – a method that has been frequently used to estimate PV cell innervation (see for example (Di Cristo et al., 2004; Feldmeyer et al., 2006; Hill et al., 2012)) – we estimated the number of somatic contacts per Pyr neuron, then compared these values using a neurite-to-puncta distance detection threshold of 0.15µm (Supplemental Figure S5).

The use of multiple features (presynaptic neurite, postsynaptic puncta, and postsynaptic neuron) to quantify input density reduced the number of false positives from non-synaptic neurite juxtaposition, and reduced the number of false negatives, where multiple discrete synapses were found beneath a contiguous region of axonal contact (Supplemental Figure S5). Using this correction, FAPpost puncta at the soma were more than 4-fold more likely to be aligned with PV than SST neurites (mean±SD, somatic PV-assigned puncta 66±30, n=9 Pyr neurons; versus somatic SST-assigned puncta 15±8, n=9 Pyr neurons, Figure 7A,G,H and Supplemental Figure S5). Analysis of VIP-associated inputs revealed a small number of colocalized post-synaptic puncta at the soma (somatic VIP-assigned puncta 11±16, <5% of total somatic puncta; n=9 Pyr neurons). Approximately one-third of somatic puncta were assigned to specific inputs; the stringency of our input-detection parameters likely underestimates the number of contacts, particularly for PV neurons. Our findings are consistent with prior reports showing that the majority of somatic inputs arise from PV neurons, with a minority of other inhibitory inputs (Di Cristo et al., 2004; Hill et al., 2012; Kubota et al., 2016; Melchitzky and Lewis, 2008; Micheva and Beaulieu, 1995), and show that synapse identification improves the accuracy of quantitative input analysis.

**Figure 7:**
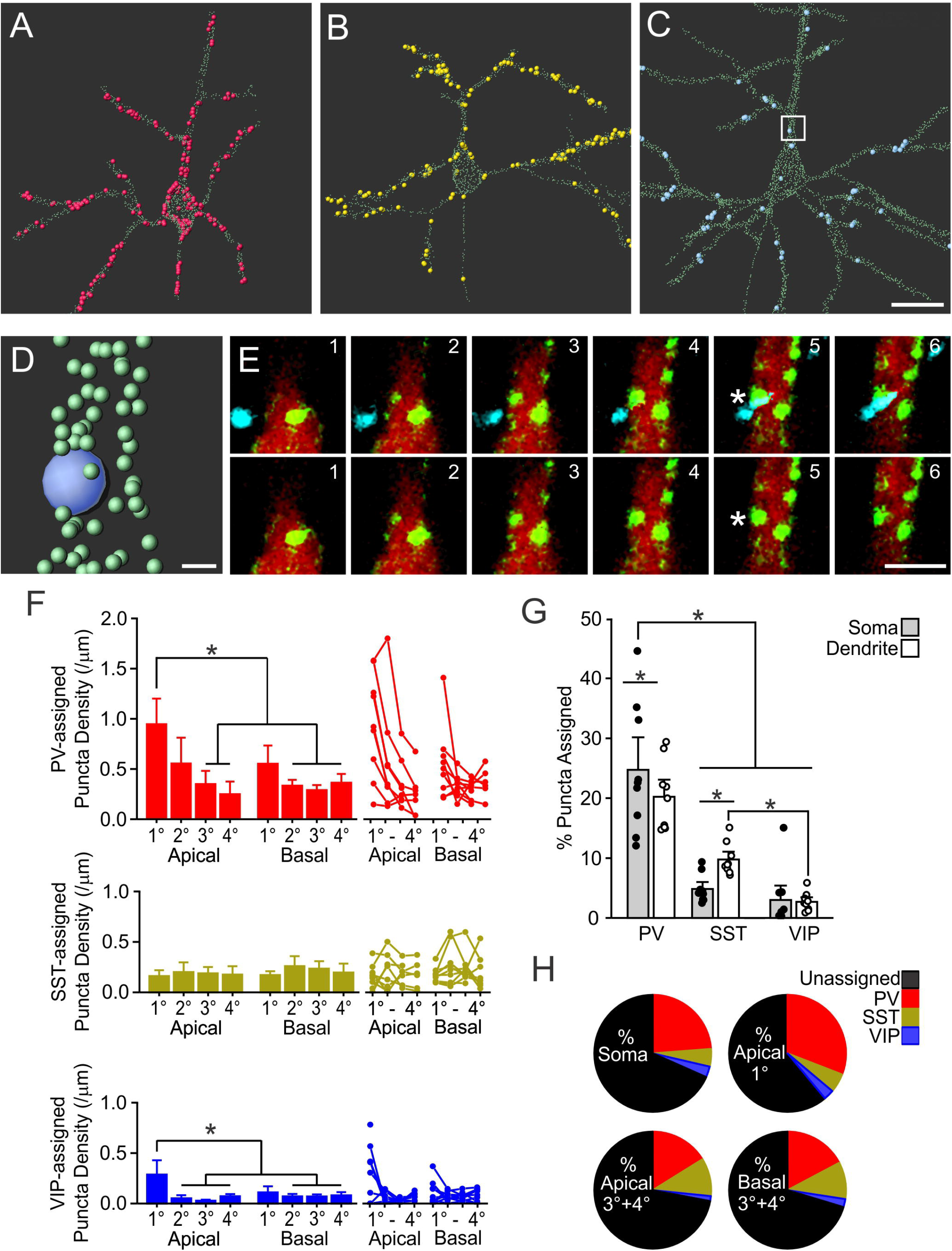
Differential distribution of inhibitory inputs across the Pyr dendritic arbor. (A) PV-input assigned synapses for an example L2 Pyr neuron. Small light-green spheres are un-assigned FAPpost puncta; large colored spheres are input-assigned FAPpost puncta. (B) As in *A* but for SST. (C) As in *A*but for VIP. Scale=20µm. See Supplemental Figure S7 for all PV, SST, and VIP input-analyzed Pyr neurons. (D) Zoom of boxed region in *C*. Scale=1µm. (E) Serial optical sections corresponding to the segment in *D* showing VIP (blue)-associated puncta (green) on the Pyr apical dendrite (red). Scale=3µm. (F) Mean density of PV, SST, and VIP-assigned FAPpost synapses across dendritic branch orders. (left) Bar is mean+SEM of all cells; (right) Individual cell values, plotted as connected lines. All data shown, statistical comparisons performed on balanced data. Reported statistics are for balanced data. PV-assigned puncta density was greater for the 1° apical dendrite. RM ANOVAPV-Input: F(7,28)=6.7, *p*=0.002; n=5, N=3. SST-assigned puncta density was not statistically significantly different across branch orders. RM ANOVASST-Input: F(7,28)=0.63, *p*=0.7; n=5, N=3. VIP-assigned puncta density was greater for the 1° apical dendrite. RM ANOVAVIP-Input: F(7,42)=3.8, *p*=0.003; n=7, N=3. (G) Inhibitory innervation of Pyr neurons, expressed as a percent of the total number of detected synapses, for each input source. All dendritic compartments pooled for comparison. Two-Way RM ANOVACompartment x Input: F(2,24)=6.0, *p*=0.008. (H) Pie-chart showing average proportion of input-assigned FAPpost synapses versus total detected synapses, binned as perisomatic (soma and 1° apical) or higher-order dendritic compartments (apical 3°+4° and basal 3°+4°). For higher-order basal branches, all input sources were significantly different (PV=17.3±1.4%, n=7, N=4; SST=10.2±1.4%, n=7, N=4; VIP=2.3±1.4%, n=7, N=2). Two-Way RM ANOVACompartment x Input: F(5.3,48)=9.2, *p*<0.001. Two-Way RM ANOVAInput: F(2,18)=47, *p*<0.001. *Tukey post-hoc pairwise comparison test, *p*<0.05. See also Supplemental Figure S5, S6, and S7.

### 3.8 Somatic and dendritic inhibition is dominated by PV input

It is commonly held that PV inputs preferentially target the soma and SST inputs, the dendrites. However, quantitative evidence for this is lacking and indeed recent reports suggest that PV inputs are broadly observed across the dendritic arbor (Kubota et al., 2015). We compared the distribution of PV, SST and VIP inputs across the reconstructed somatodendritic arbor, including 4° branches that could extend >140µm from the soma center (Figure 7).

Quantitative input assignment revealed that PV inputs were frequently observed along the dendrites, where their distribution only modestly declined at higher branch orders (up to 4° branches; Figure 7F,H). Dendritic SST inputs were less abundant than PV inputs (mean density dendritic SST-assigned puncta 0.20±0.03/µm versus PV-assigned puncta 0.38±0.07/µm, excluding the 1° apical dendrite for both). Even in higher-order (3° and 4°) apical dendrites, SST inputs were not more numerous than PV inputs (Figure 7H). It remains possible that the L1 apical tuft of L2 Pyr neurons (that was not included in our analysis, due to sectioning artifacts) may contain dense SST inputs. Overall, quantitative input analysis shows that PV inputs are the dominant source of somatic and dendritic inhibition to Pyr neurons in L2/3.

Interestingly, we observed a pronounced concentration of PV inputs at the synapse-dense 1° apical dendrite of L2 Pyr neurons (Figure 7A,F,H), with a significant 6-fold greater density than for SST inputs. These data suggest that the 1° apical compartment might be an extension of the soma with respect to PV presynaptic targeting and synaptic integration properties.

VIP inputs to Pyr neurons showed a slightly higher density for the 1° apical versus other dendrites, although the absolute number of synapses was very low. Overall, VIP input density was 10-fold lower than PV inputs to the 1° apical dendrite, a difference that was highly significant (Figure 7H). For higher-order dendrites, VIP input density was significantly lower than PV and SST inputs (mean density, dendritic VIP-assigned puncta excluding the 1° apical dendrite 0.07±0.01/µm; see also: (Zhou et al., 2017). These differences were also reflected in the proportion of cell-type specific inputs. For example, at higher-order apical dendrites the proportion of total SST-assigned puncta (10.5±1.9% of total inputs) was slightly lower than the proportion of PV-assigned puncta (16.2±1.9% of total inputs), but significantly greater than VIP inputs (1.4±1.9%).

Paired-cell recording data indicate that SST neurons do not form chemical synapses with each other (Urban-Ciecko and Barth, 2016). We used this observation to assess false positive rates (Supplemental Figure S6). Using our analysis parameters, we found that the proportion of SST-assigned puncta on SST cells was very low (2.2±2.4%). Because this number is similar to the fraction of VIP-assigned puncta on Pyr cells, it is possible that detected VIP inputs to L2 Pyr neurons are also false positives.

These data are consistent with meticulous neuroanatomical reconstructions of synaptically-connected pairs showing that PV synapses can be observed across the dendritic arbor of neocortical pyramidal neurons (Kubota et al., 2015). In addition, the prominent absence of SST inputs at the somatic and 1° apical dendrite and higher-order dendritic branches suggests that SST neurons may selectively avoid these PV-input-enriched perisomatic compartments (Figure 7H).

### 4 Discussion

Synapses are a critical determinant of neural function, and their individual and collective properties can provide insight into how brain circuits are organized and changed by experience. Electrophysiological measurements of mEPSC and mIPSCs have been widely used to assess circuit-level adaptations in synaptic function, but typically sample only a small subset of inputs onto a neuron close to the recording electrode due to electrical filtering of small and distant signals. In contrast, anatomical methods offer a highly quantitative, compartment-specific and anatomically broad view of how synapses and cell-type specific inputs are distributed onto a neuron.

A fluorescence-based, molecular genetic platform for synaptic detection and quantitation has multiple advantages for high-throughput analysis. First, the brightness of FAP/YFPpost synaptic tags enable direct visualization of synapses in both live and fixed tissue without amplification, making them accessible tools for broad scale use. Second, synaptically-targeted fluorophores can be sparsely expressed in brain tissue, not just cultured neurons, to reveal properties of synaptic and input organization in a complex neural circuit. Third, fluorescence imaging enables use of multiple, spectrally distinct channels for cell-type-selective identification of axonal inputs and specific molecules that can differentiate synapses. Fourth, volumetric data collection is rapid and requires only a confocal microscope, and images can be used for high-throughput, automated analysis. Overall, quantitative and high-throughput synapse detection with FAP/YFPpost will facilitate cell-type specific characterization of synapses and connectivity changes across multiple animals and diverse experimental conditions.

### 4.1 Synaptic quantitation without perturbation

Experimental evidence indicates that FAP/YFPpost labels both excitatory and inhibitory synapses. FAP/YFPpost puncta are localized to somatic and dendritic shafts, where inhibitory synapses are abundant, as well as on dendritic spines where excitatory synapses lie. FAP/YFPpost fluorescence overlaps with immunostaining for gephyrin and PSD-95, markers for inhibitory and excitatory synapses. Lastly, FAPpost convergence with presynaptic inputs from confirmed GABAergic neuron subtypes, specifically PV, SST, and VIP neurons, indicates association with inhibitory inputs. The FAP/YFPpost labeling of both excitatory and inhibitory synapses allows comprehensive analysis of synapse distribution from a single postsynaptic marker. Reagents that separately enable visualization of excitatory and inhibitory synapses will also be useful tools for fluorescence-based quantitative imaging (Chen et al., 2015; Gross et al., 2013), if expression levels are high enough for reliable synapse detection.

Although it has been proposed that NL-1 is specifically targeted to excitatory synapses (Song et al., 1999), NL-1 contains a conserved binding motif for the GABAergic receptor scaffolding molecule gephyrin and is sometimes observed at inhibitory synapses (Bemben et al., 2015; Tsetsenis et al., 2014). Neurexin 1β-binding to the extracellular portion of NL-1 has been shown to enhance intracellular PSD-95 interactions (Giannone et al., 2013) and its deletion in FAP/YFPpost may enable the broader distribution observed in transduced neurons. Importantly, prior studies have shown that the absence of the extracellular NL-1 region inhibits ectopic synapse formation (Chih et al., 2005), consistent with the use of this molecule as a non-invasive and accurate tag for synapse monitoring.

Does FAPpost expression alter synaptic function *in vivo*? This is a significant issue, as the electrophysiological effects of other fluorescence synapse detection reagents have not been well-investigated (Kim et al., 2011; Martell et al., 2016). Overexpression of tagged synaptic molecules leads to an increase in overall synapse number, using electrophysiological or anatomical measurements (El-Husseini et al., 2000; Gross et al., 2013). In such cases, quantitative analysis can be misleading, reflecting either a primary overexpression effect or a secondary effect of circuit-level adjustments to abnormal synaptic input. The absence of a clear electrophysiological phenotype for the FAPpost reagent suggests that overexpression of our NL-1 tagged fluorophores, in the absence of trans-synaptic interactions, may have a minimal effect on synaptic function.

### 4.2 Synapse detection accuracy

How does FAP/YFPpost synaptic quantitation compare to previous estimates of synaptic density and input organization? Synapse density has often been estimated indirectly from fluorescence images, using spines as a proxy for synapses. This is problematic, as spine detection will underestimate synapse number by excluding shaft synapses (typically inhibitory), dually-innervated spines (>10% by some estimates (Chen et al., 2012)), spines that lie within the imaging plane, and will also undercount faintly labeled, filamentous spines.

Overall synapse densities for L2 Pyr neurons revealed an overall average of ~3 synapses/µm dendrite, a density that is well within the range of prior estimates. For example, whole-cell EM reconstructions of CA1 hippocampal neurons have shown synapse densities of 0.7-7 synapses/µm, depending on location within the dendrite (Gulyas et al., 1999). Other studies analyzing spine (not synapse) density from L2 or L5 Pyr neurons in mouse S1 report between 0.4-5.1 spines/µm, where EM studies typically reveal greater spine densities (Holtmaat et al., 2005; Kasthuri et al., 2015; Villa et al., 2016a).

Fluorescence-based synaptic tags reduce multiple sources of error that can inaccurately assess total synapse number. For isolated neurons, automated puncta assignment to the parent dendrite removes the requirement that dendritic spines be visible for synapse detection, reducing false-negative rates. This rate is still non-zero, as the distance limit set for puncta assignment (1µm) will exclude puncta that lie on longer dendritic spines, which can extend >5µm in some cases (Kasthuri et al., 2015).

False-positive (non-synaptic puncta) errors are more difficult to estimate. Intracellular pools of the targeting construct were not likely to contribute, since these were digitally excluded based upon distance to the plasma membrane; however, puncta from nearby neurons may have been inadvertently misassigned to an analyzed dendritic segment. While fluorescence-based genetic methods have advantages, they are subject to variations in expression levels, both of the labeling construct and of protein trafficking to particular synapses (for example, that may have a lower NL-1 content). It remains possible that not all synapses – for example, neuromodulatory or peptidergic inputs – were uniformly labeled using this methodology. The quantitative analysis pipeline established here attempts to reconcile high-throughput analysis with variability in synapse structure, where speed and accuracy must be balanced.

### 4.3 Volumetric imaging for high-throughput synaptic input assignment

A significant advance enabled by an all-fluorescence synaptic imaging platform is the automated assignment of cell-type specific synaptic inputs with a spectrally distinct fluorophore. The tricolor (FAPpost, dTom, and presynaptic YFP) association as a criterion for synapse detection substantially reduces the false positive rate compared to brightfield microscopy methods (Hill et al., 2012; Kubota et al., 2015; Schoonover et al., 2014). We took advantage of the complete labeling of molecularly-defined inhibitory neuron populations in Cre-driver transgenic mouse lines to examine the broad-scale distribution of FAPpost labeled synaptic inputs on L2 Pyr neurons that originated from three types of GABAergic neurons.

Our analysis revealed that PV inputs predominate at somatic locations, with approximately six times as many PV as SST inputs to the cell body. These data are consistent with reports of dense PV innervation of the soma (Kubota et al., 2016) and a small fraction of somatic SST inputs (<10%) (Hill et al., 2012), and further validate FAPpost labeling as a robust method for quantitative synapse assignment. Dendritic analysis of synapse organization identified the 1° apical dendrite of Pyr neurons as a site of particularly dense PV innervation. This aspiny region of the dendrite, particularly in neocortical Pyr neurons, has been poorly studied as prior imaging methods have not been able to reliably visualize synapses in this compartment. However, whole-neuron EM reconstructions show that >90% of inputs to the apical dendrite of CA1 neurons are inhibitory (Bloss et al., 2016; Megias et al., 2001). The distinctive properties of the apical dendrite (Major et al., 2013) suggest that PV input to this region may serve as a critical filter for top-down modulation of Pyr neuron firing in the neocortex.

Quantitative analysis showed that PV neurons have more input to Pyr neuron dendrites than other neocortical inhibitory neurons. Indeed, even when excluding the densely PV-innervated 1° apical dendrite, mean dendritic input density was greater for PV than SST, and VIP inputs. Although this may be incongruous with the simplified model that contrasts soma-targeting PV and dendrite-targeting SST inputs (Chen et al., 2015; Higley, 2014; Lazarus and Huang, 2011; Pakan et al., 2016; Pi et al., 2013), prior experimental data are much less categorical than these schema suggest. For example, anatomical reconstructions from paired whole-cell recordings show that the majority of PV inputs to neocortical pyramidal neurons are located >50µm from the soma (Hill et al., 2012; Kubota et al., 2015) and abundant SST contacts can be detected at both proximal and distal dendrites (Di Cristo et al., 2004; Hill et al., 2012; Marlin et al., 2014). It remains possible that very distal dendrites, particularly in L1, have a disproportionate association of SST inputs. Based on the density of synaptic inputs, our data indicate that PV-mediated synaptic input will be the predominant source of inhibition across the somatodendritic compartments of L2 Pyr.

### 5 Conclusion

These methods open the door for large-scale anatomical imaging to examine circuit- and brain-wide changes in synapse distribution in development, learning, and disease. Future efforts should leverage volumetric imaging in cleared or expanded tissue for complete and high-resolution capture of the entire dendritic apparatus, application of additional molecular markers to distinguish different synapse types, and will employ new presynaptic constructs for improved synaptic discrimination. A critical challenge of these future possibilities will be the digital capture and storage of large anatomical datasets for computational analysis.

## Acknowledgements

We thank Megumi Matsushita and Rogan Grant for animal surgeries, Xiaobai Ren for dissociated neocortical neuron culture, Pooja Pandya and Amanda Kuhn for assistance with Imaris 3D reconstructions, Luke McDonald for assistance with histology, Joanne Steinmiller for expert animal care, and members of the Barth lab for critical comments on the manuscript. This work was supported by the Kaufman Foundation (to ALB and MPB), NIH R21 NS092019 (to MPB and ALB), NIH RF1 MH114103 (to MPB and ALB), NIH T32 NS 086749 (to DAK), Swiss National Foundation 171978 (to KZ) and the JPB Foundation (to WX), NIH 1S10RR025488-01 (to SCW), NIH 1S10RR016236-01 (to SCW), NIH 1S10RR019003-01 (to SCW), and NIH R01 NS081678 (to DBA).

## Author contributions

DAK was responsible for all aspects of the *in vivo* construct expression and quantitative synaptic analysis and writing of the manuscript. EP contributed to synaptic quantitation and input analysis. JL carried out *in vivo* imaging. KZ and WX carried out FAPpost expression and immunostaining in cultured neocortical neurons. SCW supervised immuno-EM experiments. CAT was responsible for design and generation of the FAP/YFPpost constructs. DSA contributed to the cloning of Cre-dependent AAV vectors. DBA provided FingR viruses. MPB was responsible for design and creation of the postsynaptic targeting constructs and contributed to fluorescence image analysis, and manuscript preparation. ALB was responsible for construct and experimental design, data acquisition and analysis, and writing of the manuscript.

## Conflict of interest statement

MPB is a founder and shareholder in Sharp Edge Labs, a company that licensed and is commercially utilizing the FAP-Fluorogen technology.

## Reference

Bayes, A., van de Lagemaat, L.N., Collins, M.O., Croning, M.D., Whittle, I.R., Choudhary, J.S., and Grant, S.G. (2011). Characterization of the proteome, diseases and evolution of the human postsynaptic density. Nature Neuroscience 14, 19–21.

Bemben, M.A., Shipman, S.L., Nicoll, R.A., and Roche, K.W. (2015). The cellular and molecular landscape of neuroligins. Trends Neurosci 38, 496–505.

Bloss, E.B., Cembrowski, M.S., Karsh, B., Colonell, J., Fetter, R.D., and Spruston, N. (2016). Structured Dendritic Inhibition Supports Branch-Selective Integration in CA1 Pyramidal Cells. Neuron 89, 1016–1030.

Bock, D.D., Lee, W.C., Kerlin, A.M., Andermann, M.L., Hood, G., Wetzel, A.W., Yurgenson, S., Soucy, E.R., Kim, H.S., and Reid, R.C. (2011). Network anatomy and in vivo physiology of visual cortical neurons. Nature 471, 177–182.

Briggman, K.L., Helmstaedter, M., and Denk, W. (2011). Wiring specificity in the direction-selectivity circuit of the retina. Nature 471, 183–188.

Cane, M., Maco, B., Knott, G., and Holtmaat, A. (2014). The relationship between PSD-95 clustering and spine stability in vivo. J Neurosci 34, 2075–2086.

Chandrasekaran, S., Navlakha, S., Audette, N.J., McCreary, D.D., Suhan, J., Bar-Joseph, Z., and Barth, A.L. (2015). Unbiased, High-Throughput Electron Microscopy Analysis of Experience-Dependent Synaptic Changes in the Neocortex. J Neurosci 35, 16450–16462.

Chen, J.L., Villa, K.L., Cha, J.W., So, P.T., Kubota, Y., and Nedivi, E. (2012). Clustered dynamics of inhibitory synapses and dendritic spines in the adult neocortex. Neuron 74, 361–373.

Chen, S.X., Kim, A.N., Peters, A.J., and Komiyama, T. (2015). Subtype-specific plasticity of inhibitory circuits in motor cortex during motor learning. Nature Neuroscience 18, 1109–1115.

Chih, B., Engelman, H., and Scheiffele, P. (2005). Control of excitatory and inhibitory synapse formation by neuroligins. Science 307, 1324–1328.

Chubykin, A.A., Liu, X., Comoletti, D., Tsigelny, I., Taylor, P., and Sudhof, T.C. (2005). Dissection of synapse induction by neuroligins: effect of a neuroligin mutation associated with autism. J Biol Chem 280, 22365–22374.

Deitcher, Y., Eyal, G., Kanari, L., Verhoog, M.B., Atenekeng Kahou, G.A., Mansvelder, H.D., de Kock, C.P.J., and Segev, I. (2017). Comprehensive Morpho-Electrotonic Analysis Shows 2 Distinct Classes of L2 and L3 Pyramidal Neurons in Human Temporal Cortex. Cereb Cortex 27, 5398–5414.

Di Cristo, G., Wu, C., Chattopadhyaya, B., Ango, F., Knott, G., Welker, E., Svoboda, K., and Huang, Z.J. (2004). Subcellular domain-restricted GABAergic innervation in primary visual cortex in the absence of sensory and thalamic inputs. Nature Neuroscience 7, 1184–1186.

El-Husseini, A.E., Schnell, E., Chetkovich, D.M., Nicoll, R.A., and Bredt, D.S. (2000). PSD-95 involvement in maturation of excitatory synapses. Science 290, 1364–1368.

Feldmeyer, D., Lubke, J., and Sakmann, B. (2006). Efficacy and connectivity of intracolumnar pairs of layer 2/3 pyramidal cells in the barrel cortex of juvenile rats. The Journal of Physiology 575, 583–602.

Fortin, D.A., Tillo, S.E., Yang, G., Rah, J.C., Melander, J.B., Bai, S., Soler-Cedeno, O., Qin, M., Zemelman, B.V., Guo, C., et al. (2014). Live imaging of endogenous PSD-95 using ENABLED: a conditional strategy to fluorescently label endogenous proteins. J Neurosci 34, 16698–16712.

Giannone, G., Mondin, M., Grillo-Bosch, D., Tessier, B., Saint-Michel, E., Czondor, K., Sainlos, M., Choquet, D., and Thoumine, O. (2013). Neurexin-1beta binding to neuroligin-1 triggers the preferential recruitment of PSD-95 versus gephyrin through tyrosine phosphorylation of neuroligin-1. Cell Rep 3, 1996–2007.

Glausier, J.R., Roberts, R.C., and Lewis, D.A. (2017). Ultrastructural analysis of parvalbumin synapses in human dorsolateral prefrontal cortex. J Comp Neurol 525, 2075–2089.

Gross, G.G., Junge, J.A., Mora, R.J., Kwon, H.B., Olson, C.A., Takahashi, T.T., Liman, E.R., Ellis-Davies, G.C., McGee, A.W., Sabatini, B.L., et al. (2013). Recombinant probes for visualizing endogenous synaptic proteins in living neurons. Neuron 78, 971–985.

Gulyas, A.I., Megias, M., Emri, Z., and Freund, T.F. (1999). Total number and ratio of excitatory and inhibitory synapses converging onto single interneurons of different types in the CA1 area of the rat hippocampus. J Neurosci 19, 10082–10097.

Hersch, S.M., and White, E.L. (1981). Quantification of synapses formed with apical dendrites of Golgi-impregnated pyramidal cells: variability in thalamocortical inputs, but consistency in the ratios of asymmetrical to symmetrical synapses. Neuroscience 6, 1043–1051.

Higley, M.J. (2014). Localized GABAergic inhibition of dendritic Ca(2+) signalling. Nat Rev Neurosci 15, 567–572.

Hill, S.L., Wang, Y., Riachi, I., Schurmann, F., and Markram, H. (2012). Statistical connectivity provides a sufficient foundation for specific functional connectivity in neocortical neural microcircuits. Proc Natl Acad Sci U S A 109, E2885–2894.

Hippenmeyer, S., Vrieseling, E., Sigrist, M., Portmann, T., Laengle, C., Ladle, D.R., and Arber, S. (2005). A developmental switch in the response of DRG neurons to ETS transcription factor signaling. PLoS Biol 3, e159.

Holtmaat, A.J., Trachtenberg, J.T., Wilbrecht, L., Shepherd, G.M., Zhang, X., Knott, G.W., and Svoboda, K. (2005). Transient and persistent dendritic spines in the neocortex in vivo. Neuron 45, 279–291.

Jiang, X., Shen, S., Cadwell, C.R., Berens, P., Sinz, F., Ecker, A.S., Patel, S., and Tolias, A.S. (2015). Principles of connectivity among morphologically defined cell types in adult neocortex. Science 350, aac9462.

Kasthuri, N., Hayworth, K.J., Berger, D.R., Schalek, R.L., Conchello, J.A., Knowles-Barley, S., Lee, D., Vazquez-Reina, A., Kaynig, V., Jones, T.R., et al. (2015). Saturated Reconstruction of a Volume of Neocortex. Cell 162, 648–661.

Kim, J., Zhao, T., Petralia, R.S., Yu, Y., Peng, H., Myers, E., and Magee, J.C. (2011). mGRASP enables mapping mammalian synaptic connectivity with light microscopy. Nat Methods 9, 96–102.

Kim, J.S., Greene, M.J., Zlateski, A., Lee, K., Richardson, M., Turaga, S.C., Purcaro, M., Balkam, M., Robinson, A., Behabadi, B.F., et al. (2014). Space-time wiring specificity supports direction selectivity in the retina. Nature 509, 331–336.

Kornfeld, J., Benezra, S.E., Narayanan, R.T., Svara, F., Egger, R., Oberlaender, M., Denk, W., and Long, M.A. (2017). EM connectomics reveals axonal target variation in a sequence-generating network. ELife 6.

Kremers, G.J., Goedhart, J., van Munster, E.B., and Gadella, T.W., Jr. (2006). Cyan and yellow super fluorescent proteins with improved brightness, protein folding, and FRET Forster radius. Biochemistry 45, 6570–6580.

Kubota, Y., Karube, F., Nomura, M., and Kawaguchi, Y. (2016). The Diversity of Cortical Inhibitory Synapses. Front Neural Circuits 10, 27.

Kubota, Y., Kondo, S., Nomura, M., Hatada, S., Yamaguchi, N., Mohamed, A.A., Karube, F., Lubke, J., and Kawaguchi, Y. (2015). Functional effects of distinct innervation styles of pyramidal cells by fast spiking cortical interneurons. ELife 4.

Larkum, M.E., Nevian, T., Sandler, M., Polsky, A., and Schiller, J. (2009). Synaptic integration in tuft dendrites of layer 5 pyramidal neurons: a new unifying principle. Science 325, 756–760.

Larkum, M.E., Waters, J., Sakmann, B., and Helmchen, F. (2007). Dendritic spikes in apical dendrites of neocortical layer 2/3 pyramidal neurons. J Neurosci 27, 8999–9008.

Lazarus, M.S., and Huang, Z.J. (2011). Distinct maturation profiles of perisomatic and dendritic targeting GABAergic interneurons in the mouse primary visual cortex during the critical period of ocular dominance plasticity. Journal of Neurophysiology 106, 775–787.

Major, G., Larkum, M.E., and Schiller, J. (2013). Active properties of neocortical pyramidal neuron dendrites. Annu Rev Neurosci 36, 1–24.

Marlin, J.J., and Carter, A.G. (2014). GABA-A receptor inhibition of local calcium signaling in spines and dendrites. JNeurosci 34, 15898–15911.

Martell, J.D., Yamagata, M., Deerinck, T.J., Phan, S., Kwa, C.G., Ellisman, M.H., Sanes, J.R., and Ting, A.Y. (2016). A split horseradish peroxidase for the detection of intercellular protein-protein interactions and sensitive visualization of synapses. Nat Biotechnol 34, 774–780.

Megias, M., Emri, Z., Freund, T.F., and Gulyas, A.I. (2001). Total number and distribution of inhibitory and excitatory synapses on hippocampal CA1 pyramidal cells. Neuroscience 102, 527–540.

Melchitzky, D.S., and Lewis, D.A. (2008). Dendritic-targeting GABA neurons in monkey prefrontal cortex: comparison of somatostatin- and calretinin-immunoreactive axon terminals. Synapse 62, 456–465.

Micheva, K.D., and Beaulieu, C. (1995). An anatomical substrate for experience-dependent plasticity of the rat barrel field cortex. Proc Natl Acad Sci U S A 92, 11834–11838.

Micheva, K.D., and Smith, S.J. (2007). Array tomography: a new tool for imaging the molecular architecture and ultrastructure of neural circuits. Neuron 55, 25–36.

Munoz, W., Tremblay, R., Levenstein, D., and Rudy, B. (2017). Layer-specific modulation of neocortical dendritic inhibition during active wakefulness. Science 355, 954–959.

Pakan, J.M., Lowe, S.C., Dylda, E., Keemink, S.W., Currie, S.P., Coutts, C.A., and Rochefort, N.L. (2016). Behavioral-state modulation of inhibition is context-dependent and cell type specific in mouse visual cortex. ELife 5.

Pi, H.J., Hangya, B., Kvitsiani, D., Sanders, J.I., Huang, Z.J., and Kepecs, A. (2013). Cortical interneurons that specialize in disinhibitory control. Nature 503, 521–524.

Prange, O., Wong, T.P., Gerrow, K., Wang, Y.T., and El-Husseini, A. (2004). A balance between excitatory and inhibitory synapses is controlled by PSD-95 and neuroligin. Proc Natl Acad Sci U S A 101, 13915–13920.

Pratt, C.P., Kuljis, D.A., Homanics, G.E., He, J., Kolodieznyi, D., Dudem, S., Hollywood, M.A., Barth, A.L., and Bruchez, M.P. (2017). Tagging of Endogenous BK Channels with a Fluorogen-Activating Peptide Reveals beta4-Mediated Control of Channel Clustering in Cerebellum. Front Cell Neurosci 11, 337.

Pronneke, A., Scheuer, B., Wagener, R.J., Mock, M., Witte, M., and Staiger, J.F. (2015). Characterizing VIP Neurons in the Barrel Cortex of VIPcre/tdTomato Mice Reveals Layer-Specific Differences. Cereb Cortex 25, 4854–4868.

Sassoe-Pognetto, M., Wassle, H., and Grunert, U. (1994). Glycinergic synapses in the rod pathway of the rat retina: cone bipolar cells express the alpha 1 subunit of the glycine receptor. J Neurosci 14, 5131–5146.

Scheiffele, P., Fan, J., Choih, J., Fetter, R., and Serafini, T. (2000). Neuroligin expressed in nonneuronal cells triggers presynaptic development in contacting axons. Cell 101, 657–669.

Schoonover, C.E., Tapia, J.C., Schilling, V.C., Wimmer, V., Blazeski, R., Zhang, W., Mason, C.A., and Bruno, R.M. (2014). Comparative strength and dendritic organization of thalamocortical and corticocortical synapses onto excitatory layer 4 neurons. J Neurosci 34, 6746–6758.

Shi, R., Redman, P., Ghose, D., Liu, Y., Ren, X., Ding, L.J., Liu, M., Jones, K.J., and Xu, W. (2017). Shank Proteins Differentially Regulate Synaptic Transmission. eNeuro ENEURO.0163-15.2017.

Song, J.Y., Ichtchenko, K., Sudhof, T.C., and Brose, N. (1999). Neuroligin 1 is a postsynaptic cell-adhesion molecule of excitatory synapses. Proc Natl Acad Sci U S A 96, 1100–1105.

Spruston, N. (2008). Pyramidal neurons: dendritic structure and synaptic integration. Nat Rev Neurosci 9, 206–221.

Sudhof, T.C. (2017). Synaptic Neurexin Complexes: A Molecular Code for the Logic of Neural Circuits. Cell 171, 745–769.

Szent-Gyorgyi, C., Schmidt, B.F., Creeger, Y., Fisher, G.W., Zakel, K.L., Adler, S., Fitzpatrick, J.A., Woolford, C.A., Yan, Q., Vasilev, K.V., et al. (2008). Fluorogen-activating single-chain antibodies for imaging cell surface proteins. Nat Biotechnol 26, 235–240.

Szent-Gyorgyi, C., Stanfield, R.L., Andreko, S., Dempsey, A., Ahmed, M., Capek, S., Waggoner, A.S., Wilson, I.A., and Bruchez, M.P. (2013). Malachite green mediates homodimerization of antibody VL domains to form a fluorescent ternary complex with singular symmetric interfaces. J Mol Biol 425, 4595–4613.

Tamas, G., Buhl, E.H., Lorincz, A., and Somogyi, P. (2000). Proximally targeted GABAergic synapses and gap junctions synchronize cortical interneurons. Nature Neuroscience 3, 366–371.

Tang, A.H., Chen, H., Li, T.P., Metzbower, S.R., MacGillavry, H.D., and Blanpied, T.A. (2016). A trans-synaptic nanocolumn aligns neurotransmitter release to receptors. Nature 536, 210–214.

Taniguchi, H., He, M., Wu, P., Kim, S., Paik, R., Sugino, K., Kvitsiani, D., Fu, Y., Lu, J., Lin, Y., et al. (2011). A resource of Cre driver lines for genetic targeting of GABAergic neurons in cerebral cortex. Neuron 71, 995–1013.

Tasic, B., Menon, V., Nguyen, T.N., Kim, T.K., Jarsky, T., Yao, Z., Levi, B., Gray, L.T., Sorensen, S.A., Dolbeare, T., et al. (2016). Adult mouse cortical cell taxonomy revealed by single cell transcriptomics. Nature Neuroscience 19, 335–346.

Telmer, C.A., Verma, R., Teng, H., Andreko, S., Law, L., and Bruchez, M.P. (2015). Rapid, specific, no-wash, far-red fluorogen activation in subcellular compartments by targeted fluorogen activating proteins. ACS Chem Biol 10, 1239–1246.

Tsetsenis, T., Boucard, A.A., Arac, D., Brunger, A.T., and Sudhof, T.C. (2014). Direct visualization of trans-synaptic neurexin-neuroligin interactions during synapse formation. J Neurosci 34, 15083–15096.

Tyler, W.A., Medalla, M., Guillamon-Vivancos, T., Luebke, J.I., and Haydar, T.F. (2015). Neural precursor lineages specify distinct neocortical pyramidal neuron types. J Neurosci 35, 6142–6152.

Urban-Ciecko, J., and Barth, A.L. (2016). Somatostatin-expressing neurons in cortical networks. Nat Rev Neurosci 17, 401–409.

van Aerde, K.I., and Feldmeyer, D. (2015). Morphological and physiological characterization of pyramidal neuron subtypes in rat medial prefrontal cortex. Cereb Cortex 25, 788–805.

Villa, K.L., Berry, K.P., Subramanian, J., Cha, J.W., Oh, W.C., Kwon, H.B., Kubota, Y., So, P.T., and Nedivi, E. (2016b). Inhibitory Synapses Are Repeatedly Assembled and Removed at Persistent Sites In Vivo. Neuron 89, 756–769.

Viltono, L., Patrizi, A., Fritschy, J.M., and Sassoe-Pognetto, M. (2008). Synaptogenesis in the cerebellar cortex: differential regulation of gephyrin and GABAA receptors at somatic and dendritic synapses of Purkinje cells. J Comp Neurol 508, 579–591.

Vishwanathan, A., Daie, K., Ramirez, A.D., Lichtman, J.W., Aksay, E.R.F., and Seung, H.S. (2017). Electron Microscopic Reconstruction of Functionally Identified Cells in a Neural Integrator. Curr Biol 27, 2137–2147 e2133.

Yamagata, M., and Sanes, J.R. (2012). Transgenic strategy for identifying synaptic connections in mice by fluorescence complementation (GRASP). Front Mol Neurosci 5, 18.

Yamashita, T., Pala, A., Pedrido, L., Kremer, Y., Welker, E., and Petersen, C.C. (2013). Membrane potential dynamics of neocortical projection neurons driving target-specific signals. Neuron 80, 1477–1490.

Yassin, L., Benedetti, B.L., Jouhanneau, J.S., Wen, J.A., Poulet, J.F., and Barth, A.L. (2010). An embedded subnetwork of highly active neurons in the neocortex. Neuron 68, 1043–1050.

Zaitsev, A.V., Povysheva, N.V., Gonzalez-Burgos, G., and Lewis, D.A. (2012). Electrophysiological classes of layer 2/3 pyramidal cells in monkey prefrontal cortex. Journal of Neurophysiology 108, 595–609.

Zemoura, K., Trumpler, C., and Benke, D. (2016). Lys-63-linked Ubiquitination of gamma-Aminobutyric Acid (GABA), Type B1, at Multiple Sites by the E3 Ligase Mind Bomb-2 Targets GABAB Receptors to Lysosomal Degradation. J Biol Chem 291, 21682–21693.

Zhou, X., Rickmann, M., Hafner, G., and Staiger, J.F. (2017). Subcellular Targeting of VIP Boutons in Mouse Barrel Cortex is Layer-Dependent and not Restricted to Interneurons. Cereb Cortex 27, 5353–5368.

